# Integrative structural analysis of NF45-NF90 heterodimers reveals architectural rearrangements and oligomerisation on binding dsRNA

**DOI:** 10.1101/2024.08.19.607423

**Authors:** Sophie Winterbourne, Uma Jayachandran, Juan Zou, Juri Rappsilber, Sander Granneman, Atlanta G. Cook

**Affiliations:** Institute of Quantitative Biology, Biochemistry and Biotechnology, Max Born Crescent, University of Edinburgh, Edinburgh, EH9 3BF, UK; Institute of Cell Biology, Max Born Crescent, Edinburgh EH9 3BF, UK; Bioanalytics, Institute of Biotechnology, Technische Universität Berlin, 13355, Berlin, Germany; Centre for Engineering Biology, Max Born Crescent, University of Edinburgh, Edinburgh EH9 3BF, UK

## Abstract

Complexes of nuclear factors 45 and 90 (NF45-NF90) play a multitude of roles in co- and post-transcriptional RNA processing, including regulating adenosine-to-inosine editing, cassette exon and back splicing, and splicing fidelity. NF45-NF90 complexes recognise dsRNA and, in human cells, primarily interact with inverted Alu repeats (AluIRs) that are commonly inserted into introns and other non-coding RNA regions. Intronic AluIRs of ∼300 bp can regulate splicing outcomes, such as generation of circRNAs. We examined domain reorganisation of NF45-NF90 domains on dsRNAs exceeding 50 bp to gain insight into its RNA recognition properties on longer dsRNAs. Using a combination of phylogenetic analysis, solution methods (including small angle X-ray scattering and quantitative cross-linking mass spectrometry), machine learning and negative stain electron microscopy, we generated a model of NF45-NF90 complex formation on dsRNA. Our data reveal that different interactions of NF45-NF90 complexes allow these proteins to coat long stretches of dsRNA. This property of the NF45-NF90 complex has important implications for how long, nuclear dsRNAs are recognised in the nucleus and how this might promote (co)-regulation of specific RNA splicing and editing events that shape the mammalian transcriptome.

## Introduction

Nuclear factors 45 and 90 (NF45 and NF90), also known as interleukin enhancer binding factors, ILF2 and ILF3, are essential proteins in mammalian cells that are involved in many post-transcriptional processes. This includes alternative splicing (Haque *et al*, 2023), circular RNA generation by back splicing (Li *et al*, 2017), miRNA biogenesis (Grasso *et al*, 2022; Grasso *et al*, 2020; Shang *et al*, 2022), adenosine-to-inosine (A-to-I) editing (Freund *et al*, 2020; Quinones-Valdez *et al*, 2019), ribosome biogenesis (Tafforeau *et al*, 2013; Van Nostrand *et al*, 2020; Wandrey *et al*, 2015), RNA decay (Nourreddine *et al*, 2020) and control of translation (Watson *et al*, 2020). Despite the multitude of contributions of NF45-NF90 complexes in post-transcriptional processes, there is currently no clear molecular model for their functions in cells.

The NF45-NF90 complex binds to dsRNA and UV cross-linking analysis in human cells has shown that NF90 primarily associates with Alu elements, particularly in introns and 3’UTRs (Quinones-Valdez *et al*., 2019; Van Nostrand *et al*., 2020). Alu elements are abundant in primate genomes, making up ∼10% of the human genome, with ∼50% of Alu elements mapping to intronic regions (Zhang *et al*, 2014b). As NF90 cross-links to both forward and reverse Alu elements in cells, and Alu inverted repeats (AluIRs) can form extended stretches of ∼ 300 bp dsRNA, it is likely that the primary association with transcripts is via AluIRs (Li *et al*., 2017; Quinones-Valdez *et al*., 2019). A large subset of circRNA biogenesis events are driven by complementary AluIR sequences in adjacent introns (Zhang *et al*., 2014b) and NF90 is hypothesized to stabilize these sequences to promote circRNA formation (Li *et al*., 2017). AluIRs are also promiscuously edited in the nucleus by adenine deaminases acting on RNA (ADARs), with >90% of editing sites mapping to Alu sequences (Levanon *et al*, 2004). Knockdown of NF45-NF90 increases A-to-I editing, suggesting these complexes may compete with ADARs for their substrates (Freund *et al*., 2020; Quinones-Valdez *et al*., 2019).

NF90, and its splice variant NF110, are made up of three folded domains. At the N-terminus is a “<u>d</u>omain associated with <u>z</u>inc fingers” (DZF) domain that heterodimerizes with a homologous domain in NF45 (Wolkowicz & Cook, 2012). This is followed by two <u>d</u>s<u>R</u>NA <u>b</u>inding <u>d</u>omains (dsRBD1 and dsRBD2) that are connected by a 50 amino acid linker. NF90 and NF110 splice variants have a segment of low complexity sequence after dsRBD2. These two isoforms differ in the C-terminal region after residue 688, which generates a short sequence in NF90 and a longer, low complexity sequence in NF110 (Castella *et al*, 2015; Duchange *et al*, 2000). The two splice isoforms differ in their subnuclear localisation and may have isoform-specific roles (Damianov *et al*, 2016; Viranaicken *et al*, 2011; Wandrey *et al*., 2015).

We previously showed, through structural studies, that the DZF domains in NF90 and NF45 are pseudoenzymes, i.e. the domain fold belongs to the family of nucleotidyltransferases, but key catalytic residues have been lost (Wolkowicz & Cook, 2012). This family includes enzymes such as poly(A) polymerase and key enzymes of the innate immune response including oligoadenylate synthases (OAS) and cyclic GMP-AMP synthase (cGAS) (Kuchta *et al*, 2009). OAS and cGAS have a common mechanism of activation whereby association of the nucleotidyltransferase domain with dsRNA or dsDNA, respectively, allosterically activates their catalytic activity (Civril *et al*, 2013; Donovan *et al*, 2013; Donovan *et al*, 2015; Hartmann *et al*, 2003; Lohofener *et al*, 2015; Zhang *et al*, 2014a). Binding to dsRNA by NF45-NF90 complexes is driven by the two dsRBDs but the presence of the DZF domain heterodimer increases affinity more than 10-fold, suggesting that this domain contributes to dsRNA binding, perhaps analogously to OAS and cGAS enzymes (Jayachandran *et al*, 2016). However, the DZF domain heterodimer alone does not show appreciable RNA binding activity (Wolkowicz & Cook, 2012).

While our previous studies provided insight into recognition of short segments of dsRNA by the tandem dsRBDs of NF90 (Jayachandran *et al*., 2016), how RNA is recognised in the context of the longer NF45-NF90 complex is not known. Furthermore, how the conformation of multidomain NF45-NF90 complexes alters on RNA binding is likely to be important for its molecular function, yet this is not well understood. Here we use an integrative structural biology approach, combining deep phylogenetic analysis, machine learning, solution-based biophysical methods and electron microscopy, to provide insights into architectural alterations in the NF45-NF90 complex on binding to dsRNA. These studies suggest that NF45-NF90 complexes can coat long segments of dsRNA, both by sandwiching the dsRNA between complexes and through lateral interactions along the dsRNA. This property suggests a unifying molecular mechanism for NF45-NF90 function *in vivo*.

## Materials and Methods

### Phylogenetic and missense mutation analysis of NF110, NF90 and NF45

Orthologous sequences were initially collected using the orthologous matrix browser (OMA, www.omabrowser.org) selecting OMA groups 1190687 (168 sequences, ILF2/NF45) and 1198895 (131 sequences, ILF3/NF90) (Altenhoff *et al*, 2021). These groups were supplemented with additional sequences from PSI-BLAST (Altschul *et al*, 1997) and iteratively using MAFFT (Katoh & Standley, 2013) with visual inspection in AliView (Larsson, 2014) to remove incomplete or low quality sequences.

Alignments were used as input files for ConSurf conservation analysis using full-length AlphaFold2 models of all proteins (Varadi *et al*, 2024). Conservation scores were extracted from the resulting .pdb files using Conservation_score_converter.py. Population missense mutation datasets were downloaded from gnomAD v2.1.1, filtered to remove pathogenic variants and formatted for 1D plotting (Deak & Cook, 2022). A modified version of Plot Protein (Turner, 2013) was used to generate figures combining tracks of conservation score and missense mutations plotted per residue for each protein. Domain annotations and secondary structure elements were based on solved structures (Jayachandran *et al*., 2016; Wolkowicz & Cook, 2012).

### NF45-NF90 protein complex expression and purification

Mouse NF90_long_ (residues 1 to 591) and full-length human NF45 were expressed separately in *E. coli* BL21(*DE3*) cells (Novagen) containing pRIPL plasmids (Agilent) encoding tRNAs for rare codons, in 2XTY media. NF90_long_ was expressed as a non-cleavable C-terminally 6xHis-tagged protein and NF45 as a cleavable N-terminal GST-tagged protein. Cells were induced at 20°C overnight with 0.3 mM IPTG. Cell pellets were co-lysed together in lysis buffer (40mM Tris-HCl pH 7.5, 500mM NaCl and 1 mM DTT) with 100 mM Pefabloc (Roche) and protease inhibitor cocktail (Roche cOmplete EDTA-free) using a cell disruptor (Constant Systems). The tagged proteins were extracted as a complex from clarified lysates by binding to GSH resin (Cytiva), packed into a column and eluted with lysis buffer supplemented with 20mM reduced glutathione (pH 7.5). The eluted protein complex was dialysed overnight in 40 mM Tris-HCl pH 8.0; 200 mM NaCl; 10mM imidazole (pH 8.0); 1 mM β-mercaptoethanol. Rhinovirus 3C protease was included to allow simultaneous cleavage of the GST tag from NF45. The dialysed protein complex was then bound to Ni^2+^-NTA resin (Sigma) and eluted over a 0.01 – 0.5 M imidazole gradient. Eluted protein complex was dialysed overnight into 40 mM Tris-HCl pH 7.5; 100 mM NaCl; 1 mM DTT. Heparin Sepharose chromatography (Cytiva) was used to remove nucleic acid contamination with sample eluted using a gradient from 0.01 – 1 M NaCl. Finally, the protein complex was purified using size exclusion chromatography (SEC, Superdex 200, Cytiva) in 20 mM HEPES pH 7.5; 150 mM NaCl; 1 mM DTT.

### Small angle X-ray scattering

The top and bottom RNA strands for 25 bp, 36 bp, and 54 bp (Suppl. Table 1) were annealed for 5 min at 95°C and cooled overnight to form dsRNA. NF45-NF90 protein was complexed with different lengths of dsRNA as follows: 2:1 and 4:1 molar ratio for NF45-NF90 protein:25 bp dsRNA and protein:36 bp dsRNA; 2:1, 4:1 and 6:1 molar ratio for NF45-NF90 protein:54 bp dsRNA. The samples were then incubated on ice for 30min before transferring to a sample loader for injection on to the SEC column.

SEC coupled to small angle X-ray scattering (SAXS) experiments were carried out at Diamond Light Source on the B21 beamline (Cowieson *et al*, 2020). Samples were loaded onto a S200 increase 3.2/200 (Cytiva) column in 20 mM HEPES; pH 7.5; 150 mM NaCl; 1 mM DTT. 60 µl of 5.5 mg/ml sample was loaded onto the column at a flow rate of 0.075 ml/min for NF45-NF90 complex mixed with dsRNA. 60 µl of 7.75-10 mg/ml was loaded onto the column at a flow rate of 0.1 ml/min for samples of NF45-NF90 constructs. Measurements were taken using an exposure time of 0.005 seconds. ScÅtter IV (https://bl1231.als.lbl.gov/scatter/) was used for solvent subtraction and basic analyses. Elution profiles were extracted in CHROMIXS (Panjkovich & Svergun, 2018). *Ab initio* bead models were generated with GASBORI for protein samples and DAMMIF for samples consisting of protein and RNA (Franke & Svergun, 2009; Svergun *et al*, 2001). GASBORI was run within the ATSAS 3.2.1 software suite, while DAMMIF was launched from within PRIMUS (Manalastas-Cantos *et al*, 2021). SAXS envelopes were generated in PyMOL, and then models of NF45-NF90 and A-form 25 bp, 36 bp, or 54 bp dsRNA (generated in COOT) were fitted within the envelopes (Emsley *et al*, 2010; Schrodinger, 2015).

Models of NF45-NF90_long_ samples, NF45_DZF_-NF90_DZF_, and NF90_dsRBDs_ were fitted against associated SAXS data using MultiFoXS (https://modbase.compbio.ucsf.edu/multifoxs/) (Schneidman-Duhovny *et al*, 2016). Models submitted to MultiFoXS were generated by overlaying the crystal structures 4AT7 and 5DV7 with AlphaFold2 models of NF90 (AF-Q9Z1X4) and NF45 (AF-Q9CXY6) in COOT (Emsley *et al*., 2010; Jayachandran *et al*., 2016; Jumper *et al*, 2021; Wolkowicz & Cook, 2012). Unsolved sections of the crystal structures were filled in by merging parts of the AlphaFold2 models into the structures, retaining only relevant residues in coordinate files for each construct. The coordinates were then minimised in Chimera 1.16 to avoid steric clashes (Pettersen *et al*, 2004). SAXS details are given in Suppl. Table 2.

### Mass photometry

Data collection was carried out on the Refeyn TwoMP instrument. A drop of immersion oil was placed on the lens, then a coverslip with a silicon gasket was placed on top. Contrast-to-mass calibration was carried out using Bovine Serum Albumin. 18.5 µl 20 mM HEPES, pH 7.5; 150 mM NaCl; 1 mM DTT was added to the silicon gasket, a clean area of the coverslip was located, and the focus was set. 1.5 µl of protein solution at 500 nM was added and mixed, to make up a final volume of 20 µl (final concentration 37.5 nM). Videos (30 s) were recorded using AcquireMP software followed by processing and analysis, including fitting Gaussian functions to identify peaks, using the DiscoverMP software (Refeyn Ltd).

### Multi-Angle Light Scattering

100 μl of purified NF45-NF90_long_ at a concentration of 0.36 mg/ml was injected into a Superdex 200 increase column (Cytiva) in 20 mM HEPES (pH 7.5); 150 mM NaCl; 1 mM DTT. The SEC column was coupled to a Viscotek V3580 refractive index unit and Viscotek SEC-MALS 20 (Malvern Instruments) to measure the molecular mass of the protein complex and its angular dependence of light scattering using OMNISEC software (Malvern Panalytical Ltd.).

### NF45-NF90_long_ Protein-RNA complex formation for cross-linking analysis

18 bp 2’ fluorinated - dsRNA (Suppl. Table 1, a kind gift from Frank Rigo, Ionis Pharmaceuticals) was used for titrating cross-linker concentrations with NF45-NF90_long_. RNA strands were annealed for 5 min at 95°C and cooled down gradually overnight to form 18mer dsRNA. Purified NF45-NF90_long_ was complexed with 18mer dsRNA in a molar ratio of 1:1 and 2:1 by incubating the protein and RNA on ice for 30 min and passing through size exclusion chromatography (Superdex 200, Cytiva) in 20 mM HEPES pH 7.5; 150 mM NaCl; 1 mM DTT. The protein-RNA complex from the main peak was used for cross-linking titrating assay.

A 25 bp GC-rich RNA sequence was used for quantitative crosslinking (Suppl. Table 1). RNA strands were annealed for 5 min at 95°C and cooled down gradually overnight to form the 25 bp dsRNA. Purified NF45-NF90_long_ was complexed with 25 bp dsRNA in a molar ratio of 2:1 by incubating the protein and RNA on ice for 30 min and passing through size exclusion chromatography (Superdex 200, Cytiva) in 20 mM HEPES pH 7.5; 150 mM NaCl; 1 mM DTT. The protein-RNA complex from the main peak was used for quantitative cross-linking mass spectrometry.

### Sample preparation for quantitative cross-linking mass spectrometry (CLMS)

Triplicate samples of 10 μg of NF45-NF90_long_ protein complex and NF45-NF90_long_-25 bp GC-rich (Suppl. Table 1) dsRNA complex (2:1 molar ratio) were cross-linked with 30 μg 1-ethyl-3-(3-dimethylaminopropyl) carbodiimide (EDC, Thermo Fisher Scientific, EDC: protein = 3:1 w/w) in the presence of 66 μg N-hydroxysulphosuccinimide (NHS, Thermo Fisher Scientific) on ice, in the dark for 90 min. EDC is a zero-length chemical cross-linker capable of covalently linking primary amines of lysine and the protein N-terminus and, to a lesser extent, hydroxyl groups of serine, threonine and tyrosine with carboxyl groups of aspartate or glutamate. The cross-linking reaction was quenched using a final concentration of 100 mM Tris-HCl (pH 8.0). SDS gel loading dye containing 50 mM DTT was then added to the cross-linked sample. 6 μg of cross-linked sample was loaded per lane and separated for 5 min on a 4-12% Bis-Tris gel (Invitrogen) using MOPS buffer. The gel was stained with Instant Blue (Expedeon). Stained bands were excised and washed with 50 mM ammonium bicarbonate (ABC) and 100% acetonitrile (ACN) to remove the stain. The gel pieces were reduced with 10 mM DTT and alkylated with 55 mM iodoacetamide for 20 min at room temperature. Samples were treated using 13 ng/μl trypsin (Promega) overnight at 37°C (Maiolica *et al*, 2007). The sample was acidified with 0.1% trifluoroacetic acid (TFA). Digested peptides were loaded on stage-tips containing am Empore Disk C18 filter conditioned with methanol and equilibrated with 0.1% TFA (Rappsilber *et al*, 2007). The bound peptides were treated with 80% ACN/0.1% TFA prior to analysis by liquid chromatography mass spectrometry (LC-MS).

### Quantitative Cross-linking Mass Spectrometry data collection and analysis

LC-MS/MS analysis was performed using Orbitrap Fusion Lumos (Thermo Fisher Scientific) with a “high/high” acquisition strategy. The peptide separation was carried out on an EASY-Spray column (50 cm × 75 μm i.d., PepMap C18, 2 μm particles, 100 Å pore size, Thermo Fisher Scientific). Mobile phase A consisted of water and 0.1% v/v formic acid. Mobile phase B consisted of 80% v/v acetonitrile and 0.1% v/v formic acid. Peptides were loaded at a flow rate of 0.3 μl/min and eluted at 0.2 μl/min using a linear gradient going from 2% mobile phase B to 40% mobile phase B over 109 min (each sample was analysed twice), followed by a linear increase from 40% to 95% mobile phase B in 11 min. The eluted peptides were directly introduced into the mass spectrometer. MS data were acquired in the data-dependent mode with 3 s acquisition cycle. Precursor spectra were recorded in the Orbitrap with a resolution of 120,000 and *m/z* range of 300-1700. The ions with a precursor charge state between 3+ and 8+ were isolated with a window size of 1.6 m/z and fragmented using high-energy collision dissociation (HCD) with collision energy 30. The fragmentation spectra were recorded in the Orbitrap with a resolution of 15,000. Dynamic exclusion was enabled with single repeat count and 60s exclusion duration. The mass spectrometric raw files were processed into peak lists using ProteoWizard version 3.0 (Kessner *et al*, 2008), and cross-linked peptides were matched to spectra using Xi software version 1.6.751 (Mendes *et al*, 2019) with in-search assignment of monoisotopic peaks (Lenz *et al*, 2018). Search parameters were MS accuracy, 3 ppm; MS/MS accuracy, 10ppm; enzyme, trypsin; cross-linker, EDC; max missed cleavages, 4; missing mono-isotopic peaks, 2; fixed modification, carbamidomethylation on cysteine; variable modifications, oxidation on methionine; fragments, b and y ions with loss of H_2_O, NH_3_ and CH_3_SOH.

Label free quantitation on MS1 level was performed using Skyline version 19.1 (MacLean *et al*, 2010). Autovalidated cross-linked peptides were introduced as an .ssl file following the standard format for custom spectral libraries in Skyline. In the .ssl file, an entry is generated for each cross-linking feature. A cross-linking feature is defined as a unique PSM for a cross-linked peptide with differences in charge state, linkage sites, or modification (Muller *et al*, 2018). Data from Skyline was exported into a .csv file for further processing in Excel to categories including control enriched (cross-linking pairs intensity higher in the control group), RNA enriched (cross-linking pairs intensity higher in the control group) and common (cross-linking pairs intensity similar in both groups)

### *In vitro* transcription

pRRG260-Cyp1A1 was a gift from Phillip Newmark (Addgene plasmid # 99081; http://n2t.net/addgene:99081;RRID:Addgene_99081) (Roberts-Galbraith & Newmark, 2013). The length of the Cyp1A1 insert was reduced by site-directed mutagenesis by designing primers so that 310 bp and 410 bp lengths of dsRNA could be generated by *in vitro* transcription (Suppl. Table 1). 50 nM of the pRRG260 Cyp1A1 plasmid mutants containing 310 bp and 410 bp were linearized using BamHI-HF (NEB) to create a run-off transcription template. *In vitro* transcription was performed using T7 RNA polymerase and NTPs. Pyrophosphatase was added to reduce inorganic pyrophosphate precipitation during the reaction. After 3 hrs incubation at 37°C, DNase I was added and incubated further at 37°C for 30 min to remove the template. The *in vitro* transcribed dsRNA was purified by passing through a PD spintrap G-25 column with a bed volume 0.6ml (GE).

### pyRBDome analysis

To generate RNA/ligand-binding site predictions for NF90 and NF45, the AlphaFold2 structures (AF-Q12906-F1 and AF-Q12905-F1 respectively) were submitted to the pyRBDome pipeline (Chu *et al*, 2024). Predictions were downloaded for the prediction algorithms BindUp (Paz *et al*, 2016), FTMap (Kozakov *et al*, 2015), RNABindRPlus (Walia *et al*, 2014), DisoRDPbind (Peng *et al*, 2017), and PST-PRNA (Li & Liu, 2022). The eXtreme Gradient Boosting ensemble models used the prediction results to calculate the probability of RNA-binding for each amino acid in the proteins. These scores were added to the B-factor column of the associated PDB file and visualised using PyMol (Schrodinger, 2015).

### Negative stain electron microscopy

NF45-NF90_long_ protein was complexed with different lengths of *in vitro*-transcribed Cyp1A1 dsRNA (310 bp and 410 bp) using a 18:1 molar ratio for NF45-NF90 protein:310 bp dsRNA and a 24:1 molar ratio for NF45-NF90 protein:410bp dsRNA. The protein and RNA were incubated for 45 min on ice. Protein-RNA samples of 4 μl (10 μg/ml) were spotted on glow discharged grids (carbon support film - 400 mesh grid, TAAB laboratories) by incubating for 2 min. The grids were then washed twice with 20 mM HEPES pH 7.5; 150 mM NaCl; 1 mM DTT and stained with 2% uranyl acetate for 2 min. Images were taken at 40X magnification using JEOL JEM-1400 Plus TEM. Representative images were collected on a GATAN OneView camera.

## Results

### Population variant analysis supports functional importance of DZF domains

To better understand the functional contribution of domains and motifs in the DZF domain proteins, we undertook an extensive phylogenetic analysis of NF45 and the two splice isoforms of the ILF3 gene, NF110 and NF90 (Fig. 1). NF90 differs from NF110 by a second splice isoform that alters the C-terminal region after residue 687 in the reference sequence (Fig. 1A). Both NF90 and NF110 have additional isoforms that include a four amino acid sequence (NVKQ) inserted after residue 516. We assembled multiple sequence alignments of both ILF3 isoforms and used the alignments, along with AlphaFold2 models, to calculate per-residue conservation scores in ConSurf (Yariv *et al*, 2023). These scores were displayed along the primary structure of all proteins (Fig. 1).

**Figure 1.**
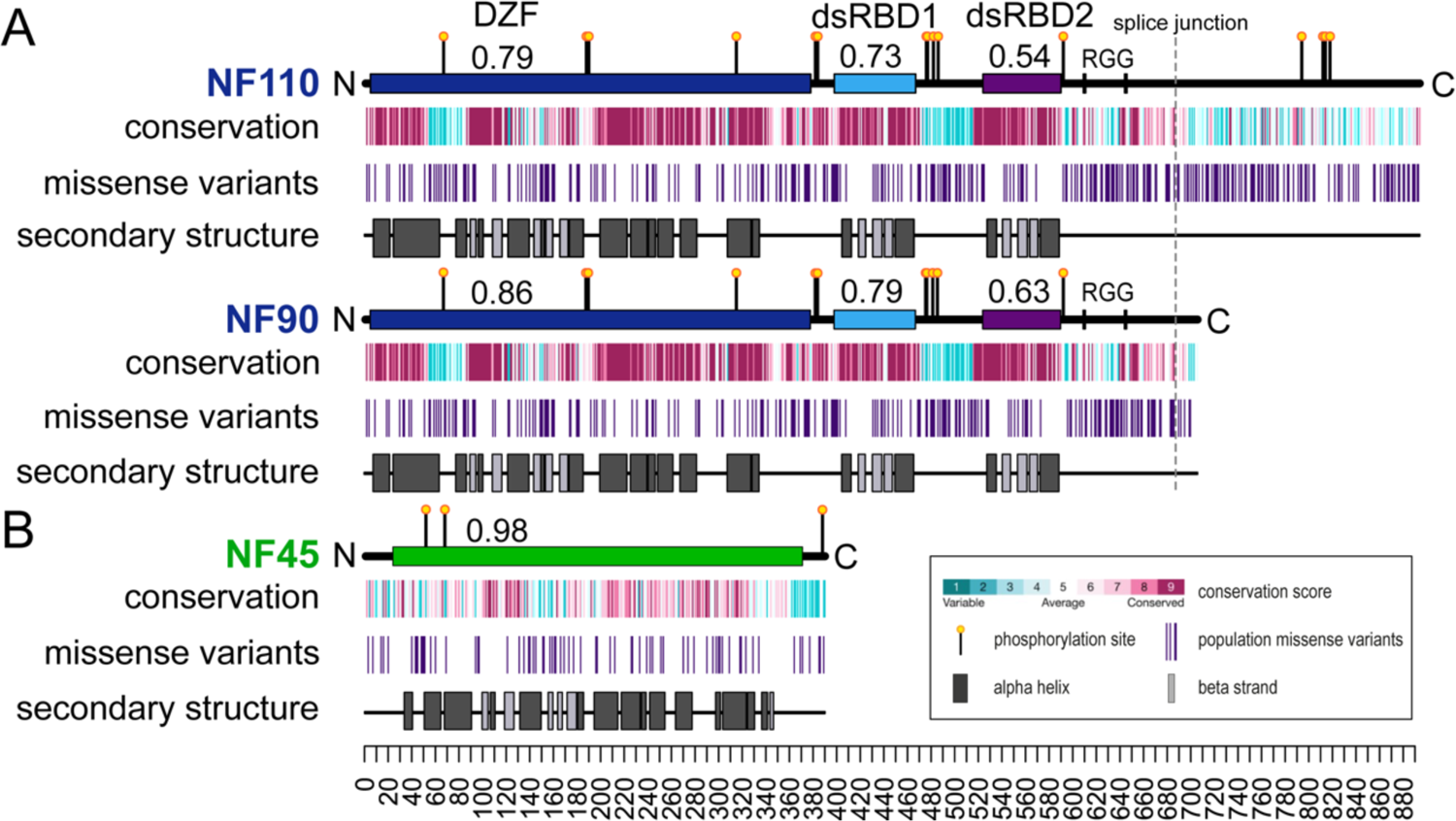
Folded domains and functional sites are conserved and depleted of benign population variants. (**A**) Primary structures of NF110 and NF90. Folded domains are indicated by dark blue, sky blue and purple rectangles along a horizontal line representing the length of the protein from N to C terminus. Fractional values associated with domains are missense depletion per domain over depletion over the whole protein (Vd/Vp) ratios. Yellow lollipops are phosphorylation sites, RGG sequences are marked with a vertical line. Three tracks show conservation (from ConSurf, where maroon is high conservation and cyan is high variation); the positions of population missense variants extracted from gnomAD (vertical lines, indigo); and positions of secondary structure elements as observed in published structures. (**B**) Similar analysis with equivalent tracks for NF45, which has only one domain (green).

We also considered population missense variants to gain additional data for domain-level functional importance (Deak & Cook, 2022). The gnomAD database contains genomic and exomic data from both healthy adults and patient data reported in ClinVar (Karczewski *et al*, 2020; Landrum *et al*, 2020; Landrum *et al*, 2014). Based on the “probability loss-of-function observed over expected upper bound” metric (pLOEUF) in gnomAD, the ILF2 and ILF3 genes are under purifying selection (0.25 and 0.1 respectively) (Karczewski *et al*., 2020). This agrees with DepMap (https://depmap.org/portal) annotations of genes ILF2 and ILF3 as common essential genes (Tsherniak *et al*, 2017). Consequently, patterns of benign missense variants of these proteins across the human population are likely to be informative on functionally important domains (Iqbal *et al*, 2020). We generated tracks for population missense variants, after removing likely pathogenic and ClinVar mutations, and compared these with the conservation scores on the primary structures of the DZF proteins (Fig. 1) (Deak & Cook, 2022). Domains and motifs with high functional importance are expected to be both conserved and locally depleted of missense variants. Calculated depletion scores (Vd/Vp ratios), where scores lower than 1 indicate localised depletion within a domain, show that the DZF domain is depleted of missense variants (Vd/Vp=0.79-0.86) compared to the rest of the protein (Deak & Cook, 2022). The dsRBD domains are both highly conserved and depleted of missense variants (Vd/Vp range 0.54-0.79) (Fig. 1), consistent with their role in RNA recognition.

The variant and conservation analyses further revealed that low complexity and/or natively unstructured regions are less well conserved than structured domains. For example, residues 57-88 in NF110/NF90 have low conservation and correspond to a region of the protein that could not be modelled in the crystal structure (Wolkowicz & Cook, 2012). Similarly, the connecting sequence between dsRBD1 and dsRBD2 of NF90 (Fig. 1A) and a glutamate-rich sequence at the C-terminus of NF45 (Fig. 1B) had low conservation scores; these also correspond to protein sequences that could not be modelled in crystallographic data (Jayachandran *et al*., 2016; Wolkowicz & Cook, 2012). As expected, these segments are correspondingly enriched in population missense variants. Overall, the patterns of conservation and population variation indicate that both the DZF domains and the dsRBD domains make significant contributions to protein function.

### NF45-NF90_long_ shows a compact architecture in solution

As NF45-NF90_long_ is a multidomain complex, understanding its architecture requires a combination of methods. We used size exclusion chromatography (SEC) coupled to small angle X-ray scattering (SAXS) to characterise the solution structure of the complex in comparison with fragments of the complex that had previously been crystallised: NF45_DZF_-NF90_DZF_ (Wolkowicz & Cook, 2012), and NF90_dsRBDs_ (Jayachandran *et al*., 2016) (Fig. 2, Fig. S1, Suppl. Table 2). NF45_DZF_-NF90_DZF_ separated as a single peak on SEC and showed no appreciable low angle scatter from aggregates by SAXS (Fig. 2B-C). Scattering curves from the peak of the SEC curve (Fig. S1A) were averaged and real space P(r) functions were calculated (Fig 2D). The maximum dimension (D_max_) derived from the NF45_DZF_-NF90_DZF_ P(r) curve was an excellent match for the longest distance measured in the X-ray crystal structure (126Å vs 124Å, respectively) (Fig. 2E). This is consistent with previous observations that the NF45_DZF_-NF90_DZF_ heterodimer is monomeric in solution (Wolkowicz & Cook, 2012). Dimensionless Kratky analysis (Fig. S1C) (Putnam *et al*, 2007; Rambo & Tainer, 2011) and *de novo* bead modelling, calculated using GASBORI (Fig. 2E) (Svergun *et al*., 2001), indicates that the NF45_DZF_-NF90_DZF_ dimer is an elongated rigid structure, consistent with the crystal structure (Fig. 2E). A similar analysis for NF90_dsRBDs_ (Fig. 2A), showed near-ideal behaviour by SEC-SAXS (Fig. 2B-C, Fig. S1A). The D_max_ value derived from the P(r) function for NF90_dsRBDs_ is 126 Å (Fig. 2D). This is consistent with the dsRBDs being connected by a flexible linker (Fig. 1A). This model is further supported by Kratky analysis, which rises at higher sR_g_ values, compared with NF45_DZF_-NF90_DZF_ (Fig. S1C) (Putnam *et al*., 2007; Rambo & Tainer, 2011), and a *de novo* bead model (Fig. 2E), consistent with a dynamic structure.

**Figure 2.**
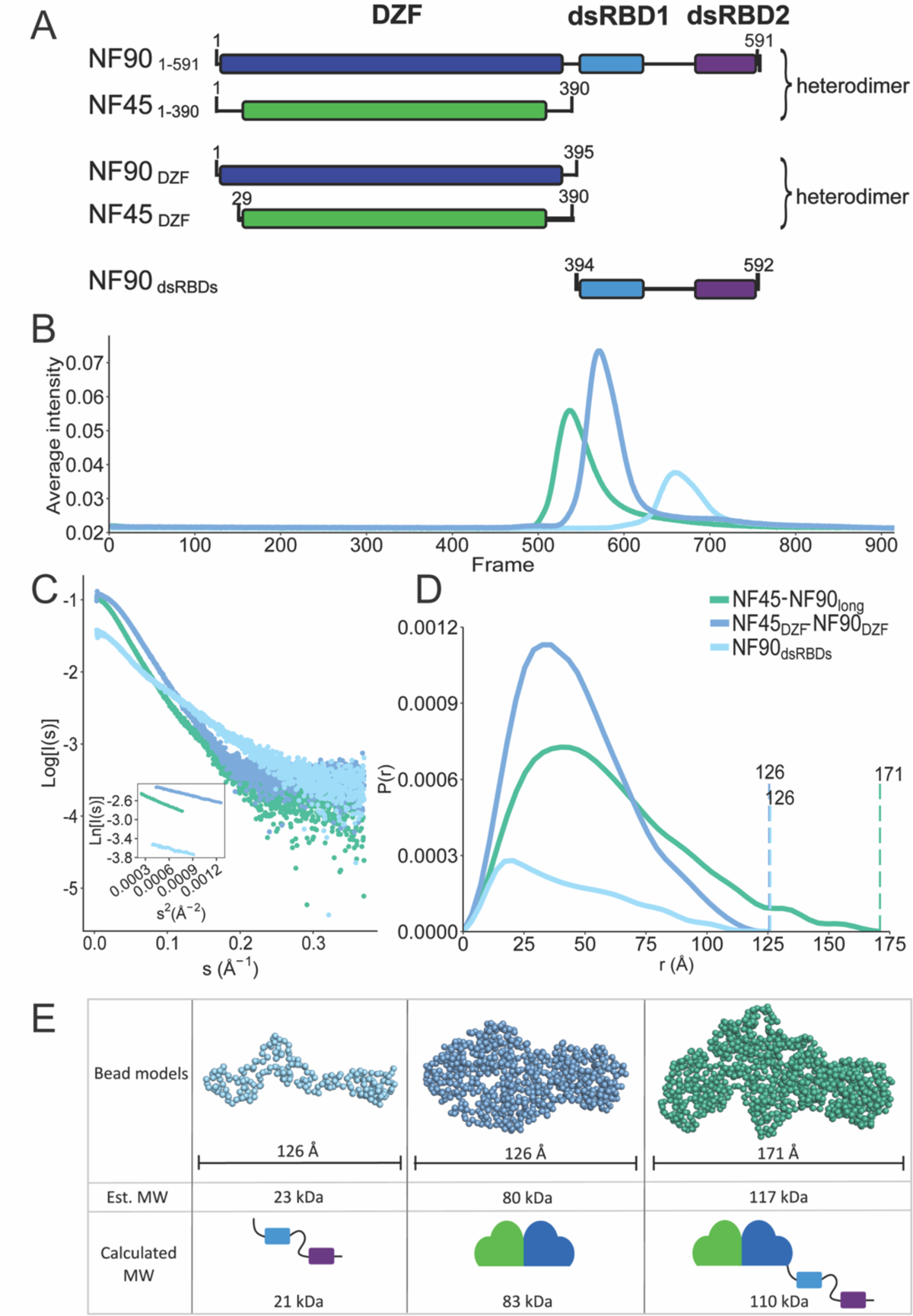
NF45-NF90 complexes in solution show compaction of domains. (**A**) Overview of constructs used for structural analyses. (**B**) SEC-SAXS profiles of three constructs. (**C**) Scattering curves and Guinier analysis (inset) of samples from (**B**), (**D**) P(r) functions derived from (**C**), (**E**) Bead models for NF45-NF90 constructs with D_max_ and calculated MW values (from amino acid composition) or estimated MW from SAXS analysis.

To understand how domain conformations compare in the context of NF45-NF90_long_, we also characterised this complex using SEC-SAXS and SEC coupled to multi-angle light scattering (MALS) (Fig. 2A-B, Fig. S1A-B). Scattering curves (Fig. 2C) were averaged from images from the middle of the SEC profile (Fig. S1A). The P(r) function derived from these data gave a D_max_ of 171 Å. Given that NF45_DZF_-NF90_DZF_ is 124 Å at its longest point and that the D_max_ of NF90_dsRBDs_ is 126 Å, this suggests that NF45-NF90_long_ is a more compact structure than expected by the combination of the three domains. This is further supported by *de novo* bead models of NF45-NF90_long_ (Fig. 2E) and Kratky analysis, which shows that NF45-NF90_long_ has a profile that is more closely related to NF45_DZF_-NF90_DZF_ than NF90_dsRBDs,_ i.e. has limited dynamics. Of the three samples characterised by SAXS, the NF45-NF90_long_ complex has the least agreement between the estimated versus the calculated molecular mass (122 kDa vs 110 kDa, respectively). A similar discrepancy is observed with SEC-MALS, where the measured molecular weight (MW) is 128 kDa (Fig. S1B). The slightly higher-than-expected MW for NF45-NF90_long_ could be explained by a small contribution of dimerization of NF45-NF90_long_ complexes.

To further explore the conformational space accessible to NF45-NF90 complexes, we used MultiFoXS (Schneidman-Duhovny *et al*., 2016) to estimate the contributions of different states of NF45-NF90_long_ in solution. All-atom models were generated for each sample, based on our prior crystal structures and AlphaFold2 models (Jumper *et al*., 2021). We searched for the best agreement with the experimental data described by the fewest possible models (Fig. S1D-G). Fitting NF45-NF90_long_ models to two independent datasets produced good fits with four or five different conformations (χ2=1.18-1.23) (Fig. S1D-E). Both sets of models included substantial contributions from structures where dsRBD1 is close to the core NF45-NF90 heterodimerisation domain, leading to a compact conformation. The NF45_DZF_-NF90_DZF_ was well described with a single model (χ^2^=1.23) (Fig. S1F), consistent with the crystal structure. As expected from *de novo* bead modelling and Kratky analysis, NF90_dsRBDs_ showed major contributions from four distinct conformations, driven by changes in linker sequence connecting the two dsRBDs (χ^2^=1.04 of fit back calculated to SAXS curve) (Fig. S1G) (Svergun *et al*., 2001). The conformational analysis further indicates that the NF90_dsRBDs_ construct accesses a wider range of conformations than the equivalent residues in a context where the DZF heterodimerisation domain is present.

### NF45-NF90_long_ interactions on dsRNA show steric exclusion

We previously showed that NF90_dsRBDs_ bind to dsRNA of a minimal length of 18 bp and that two molecules of protein typically bind to RNA (Jayachandran *et al*., 2016). Moreover, our previous studies also showed that NF45-NF90_long_ complexes have a ∼10-fold higher affinity for dsRNA than NF90_dsRBDs_ suggesting that the DZF heterodimerisation domain contributes to this interaction. To further explore binding of NF45-NF90_long_ on dsRNA, we carried out a series of SEC-SAXS measurements of NF45-NF90_long_ with dsRNA lengths of 25 bp, 36 bp and 54 bp. Comparing SEC-SAXS profiles showed that NF45-NF90_long_ alone elutes later than RNA-bound complexes, with the 54 bp dsRNA-associated complexes eluting earliest (Fig. 3A). These observations are consistent with the formation of higher order oligomers of NF45-NF90_long_ on dsRNA. Notably, some samples showed a shift to earlier elution volumes when higher protein:RNA molar ratios were used. For example, 4:1 NF45-NF90_long_:25 bp dsRNA elutes in two distinct peaks, while 2:1 NF45-NF90_long_:25 bp dsRNA shifted intensity to the earlier peak (Fig. 3A). This suggests that in the 4:1 protein:RNA samples, the RNA binding sites are saturated and excess protein elutes later.

**Figure 3.**
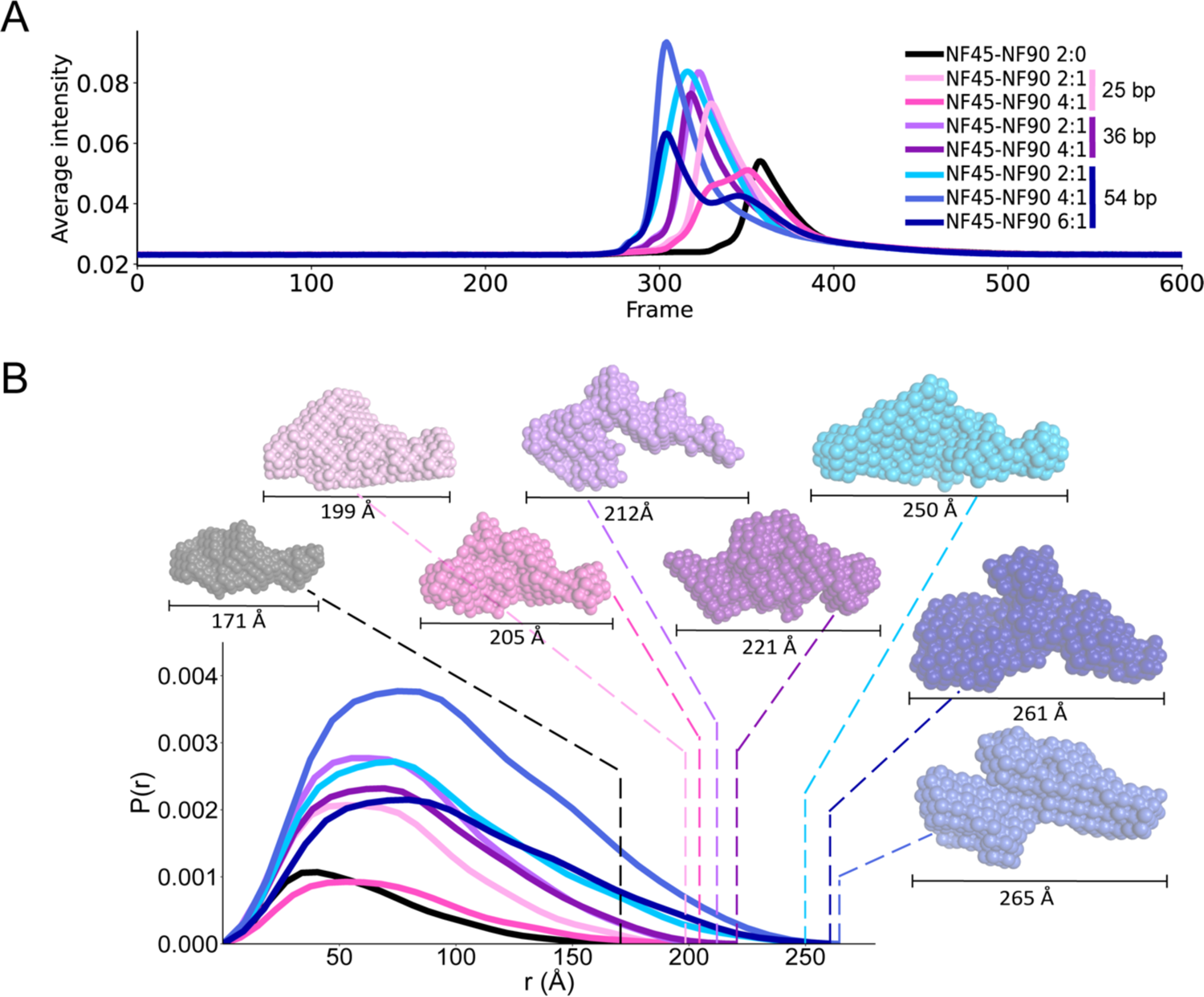
Solution analysis of NF45-NF90 binding to dsRNA of increasing lengths. Eight SAXS measurements were carried out with no dsRNA or with 25 bp, 36 bp and 54 bp RNA at increasing molar ratios of protein:dsRNA. (**A**) Intensity profiles of samples as eluted from size exclusion chromatography. Sample identities for all graphs are given in the inset key (**B**) Real space profiles of all SAXS samples indicated calculated D_max_ with dotted lines and associated bead models. D_max_ values are indicated under each model.

Early SEC peak fractions (Fig. 3A, S2A) were used to generate SAXS curves and Guinier analyses were performed (Fig. S2B). Details of analyses are given in Suppl. Table 2. Based on these data, P(r) functions were generated to extract D_max_ values for each sample (Fig. 3B) and Kratky plots were generated (Fig. S2C). Kratky analyses indicate that all protein and protein:RNA complexes are consistent with compact multi-domain structures, further suggesting that NF45-NF90_long_ complexes become more ordered on binding dsRNA. Consistent with the SEC profiles, the D_max_ values increased with increasing length of dsRNA and higher protein:RNA molar ratios. The molecular mass of each complex was estimated, and DAMMIF bead models were generated for each sample (Fig. 3B, Fig. S2D) (Franke & Svergun, 2009). These data suggest that, as expected, the number of NF45-NF90_long_ dimers bound to dsRNA increases with increasing length. However, while the minimal binding site is 18 bp, we did not observe more than 4 protein complexes per dsRNA on 54 bp. SAXS samples were characterised in parallel by mass photometry (Fig. S3) (Young *et al*, 2018), which gives the population of different complexes found in the sample. Mass photometry experiments are necessarily carried out in the range of 10-50 nM protein (i.e. ∼ 1000x lower than samples used for SEC-SAXS), such that the equilibrium for these interactions shift towards dissociation. Nevertheless, NF45-NF90_long_ complexes on 54 bp dsRNA showed distinct stoichiometries of up to 3 proteins per dsRNA (Fig. S3). This suggests that pairs of NF45-NF90_long_ complexes binding to dsRNA impose a steric constraint that extends the binding site to around 19-26 bp (Fig. S2D).

### NF45-NF90_long_ domains are rearranged on RNA binding

To further assess the effect of RNA binding on the architecture of NF45-NF90_long_ complexes, we used cross-linking and mass spectrometry (CLMS) with 1-ethyl-3-(3-dimethylaminopropyl) carbodiimide (EDC), a zero-length cross-linker (Fig. 4). Titrating EDC concentration with protein in the absence and presence of an 18 bp dsRNA, gave different patterns of bands retarded on SDS-PAGE (Fig. S4A) (Jayachandran *et al*., 2016). For protein-only samples, EDC generated two main species, one likely representing single NF45-NF90_long_ complexes, and an upper band likely representing interacting NF45-NF90_long_ complexes. Similar band patterns were present when dsRNA was bound, along with further upper bands indicating formation of larger oligomers on RNA. Retardation patterns for NF45-NF90_long_ in 1:1 protein:RNA mixtures gave a less homogeneous distribution of bands (Fig. S4A). Consequently, we generated EDC cross-link maps for NF45-NF90_long_ alone and with 18 bp dsRNA in a 2:1 protein to dsRNA ratio (Fig. 4A,B, Fig. S4A).

**Figure 4.**
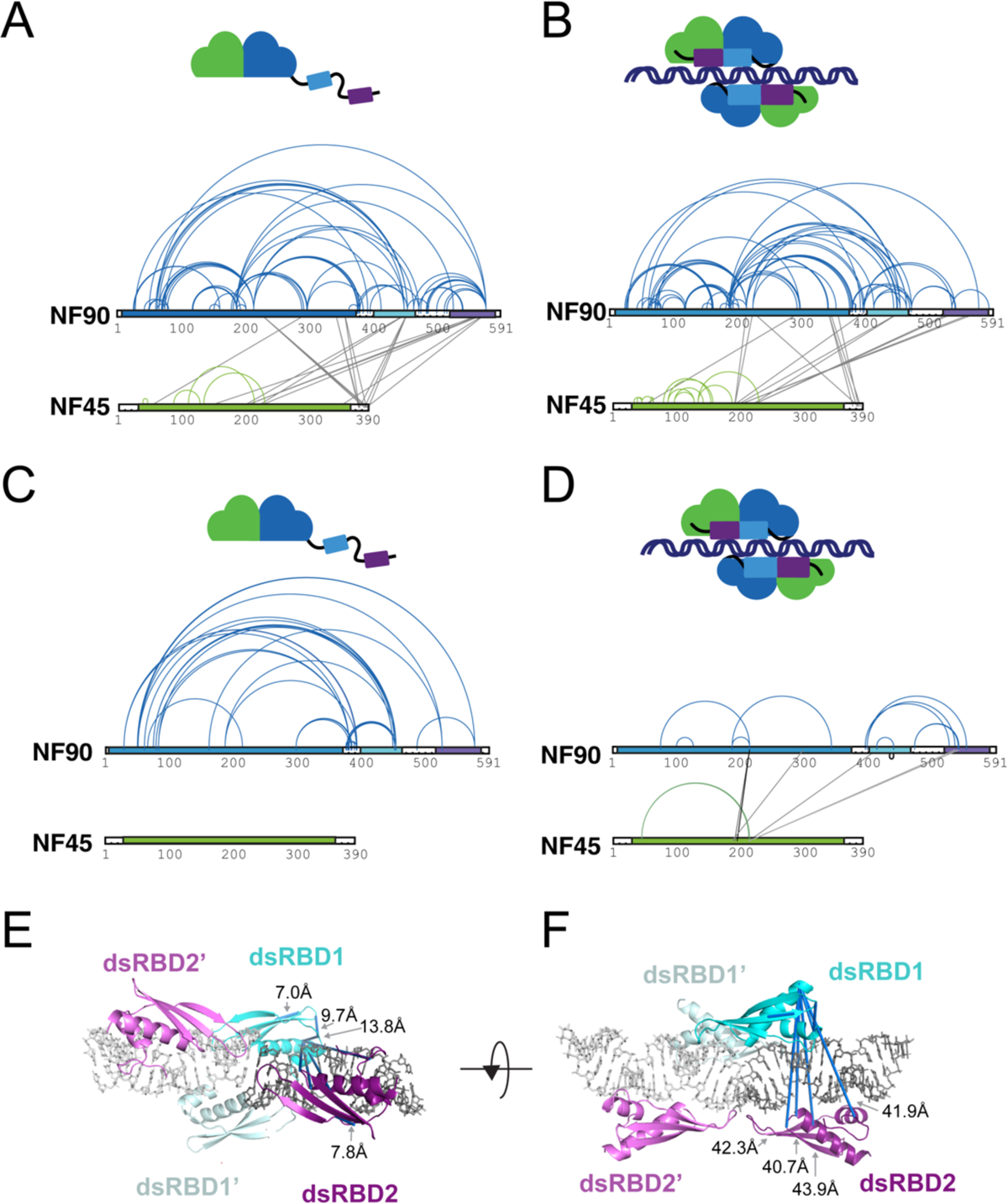
NF45-NF90 complexes undergo conformational change on binding to dsRNA. Cross-linking mass spectrometry on NF45-NF90 complexes in solution without (**A**), and with (**B**) dsRNA. Intramolecular crosslinks are blue and green for NF90 and NF45, respectively. Intermolecular crosslinks are grey. Domain limits and colouring as for Fig. 1. Quantitative differences in crosslinking comparing free NF45-NF90 (**C**) and dsRNA-bound NF45-NF90 (**D**). Cross-links with at least 2-fold differences are shown while unquantified cross-links are not displayed. (**E**) Mapping of crosslinks on to NF90d_sRBDs_. The asymmetric unit of the crystal structure is shown along with the closest symmetry equivalent of the NF90-dsRNA complex. DsRBD domains are cyan and purple with symmetry mates in lighter colours. dsRNA is shown in grey and as sticks. Shortest possible cross-links are shown in dark blue and Cα-Cα distances <24 Å are annotated. (**F**) a rotated view of the same structure is shown. Cα-Cα distances >24 Å are annotated.

Clusters of cross-links differ between samples of NF45-NF90_long_ without and with dsRNA (compare Fig. 4A and 4B), indicating that structural rearrangements have occurred. For example, a cluster of cross-links emanating from residue 577 in NF90 shows fewer links when RNA is bound. Similarly, cross-links between the C terminus of NF45 and dsRBD1 and dsRBD2 of NF90 in the RNA-free sample (Fig. 4A) are not evident when RNA is bound (Fig. 4B). The changes observed between the unbound and RNA bound samples likely reflect conformational rearrangements of dsRBD1 and dsRBD2 on binding RNA.

### Quantification of cross-links reveals RNA-dependent conformational changes

The cross-linking maps likely include contributions from unbound NF45-NF90_long_ complexes and various stoichiometries of RNA-bound states that are in equilibrium. To focus on differences specific to the unbound and RNA-bound states of NF45-NF90_long_, we repeated EDC CLMS measurements with three independent samples of unbound and RNA-bound protein using a longer, 25 bp GC-rich dsRNA. We then quantified fold changes in peak area between samples. The crosslinking maps were similar to those observed using 18 bp dsRNA (Fig. S4B). We then quantified and visualised cross-linked peptides that show a >2-fold increase or reduction between the two states (Fig. 4C,D, Suppl. Table 3), which are a fraction of the total cross-links identified across these six samples (Fig. S4B,C). The pattern of changes is consistent with changes seen with non-quantified data. For example, cross-links at the C-termini of both NF45 and NF90_long_ are reduced in intensity when RNA is bound (Fig. 4C,D).

The data imply an altered conformation of the linker region (residues 377-390) between NF90 DZF domain and dsRBD1 on RNA binding. These include residues D377, E379, and E380 which lose interactions with K454 and T452, both on dsRBD1 (Fig. 4C). Similarly, several cross-links with residues D377, E379, E380, K381, and K392 in this linker show dramatically decreased intramolecular interaction with loop 57-88 (residues E52 E75, E83) (Fig. 4C,D) when RNA is bound. The linker region is highly conserved and contains a positively charged nuclear localisation signal. It is possible that the reorientation of dsRBD1, or direct interactions with dsRNA, or both, contribute to the loss of cross-links on RNA binding. As several residues in this loop have more than one cross-linking partner, these lost interactions likely represent a shift to fewer conformational states.

Both CLMS datasets are consistent with a higher conformational freedom of the dsRBDs, particularly dsRBD2, in the absence of RNA. This is evidenced by the starkly altered pattern of interactions between the dsRBD domains and the DZF domain of NF90 when RNA is bound (Fig. 4C,D). Moreover, new inter-dsRBD interactions appear on binding RNA (D398^NF90^ to K526^NF90^, K535^NF90^, K553^NF90^; and E472^NF90^ to K540^NF90^, Y541^NF90^), as do new interactions with the DZF domain of NF45 (K413^NF90^, K535^NF90^ and K540^NF90^ to E212^NF45^ and/or E213^NF45^) (Fig. 4D). There is a corresponding loss of the intramolecular interactions of K454^NF90^ with residues E51, E61, E75, E83, and E187 on NF90 that are observed in the unbound state. Note that residues 57-88 of NF90 were not resolved in the crystal structure (Wolkowicz & Cook, 2012) and these residues lie in an unstructured, poorly conserved loop of NF90 (Fig. 1A). This further suggests ordering of dsRBD1 on binding to RNA.

Our previous structural studies of NF90_dsRBDs_ on a dsRNA substrate showed a positioning of dsRBD1 and dsRBD2 on either side of the RNA helix (Jayachandran *et al*., 2016). Moreover, a symmetry operation showed that an additional dsRBD1 and dsRBD2 could bind in proximity on a continuous dsRNA helix (Fig. 4E). To test whether the organisation of dsRBDs observed in the crystal structure match that in NF45-NF90_long_ with RNA in solution, we examined the distances of cross-links in these structural models, searching for the shortest possible distances (Fig. 4E). While cross-links within individual dsRBDs are all short (Cα-Cα 7.0Å-13.8Å) and in the expected range for EDC cross-links (<24Å), the remaining distances violate the upper limit of expected cross-link size and are ∼40-44 Å in this model. These cross-links connect residues within folded domains that are placed on opposing surfaces of the dsRNA in the solved structure (Fig. 4F) and these distances cannot be explained by local conformational flexibility of the protein chain. This analysis shows that the relative orientation of dsRBDs in the NF45-NF90_long_ complex on RNA is likely to be different to that observed in the previous crystal structures.

### Oligomers of NF45-NF90_long_ are observed on dsRNA

Among the new cross-links observed in the RNA-bound state (Fig. 4D), three cross-links show >2-fold increase in the quantified data, namely D194^NF45^-K214^NF90^, E190^NF45^-K214^NF90^ and K186^NF45^-E298^NF90^ (Fig. 4D, Fig. S4C). Mapping these cross-links onto the structure of NF45_DZF_-NF90_DZF_ heterodimers shows that these links are not possible in the context of a single NF45-NF90 complex as the distances are much longer (67-71 Å) than is possible with the EDC cross-linking chemistry (Fig. 5A). However, these cross-links could be rationalised if they occur between NF45-NF90 heterodimers.

**Figure 5.**
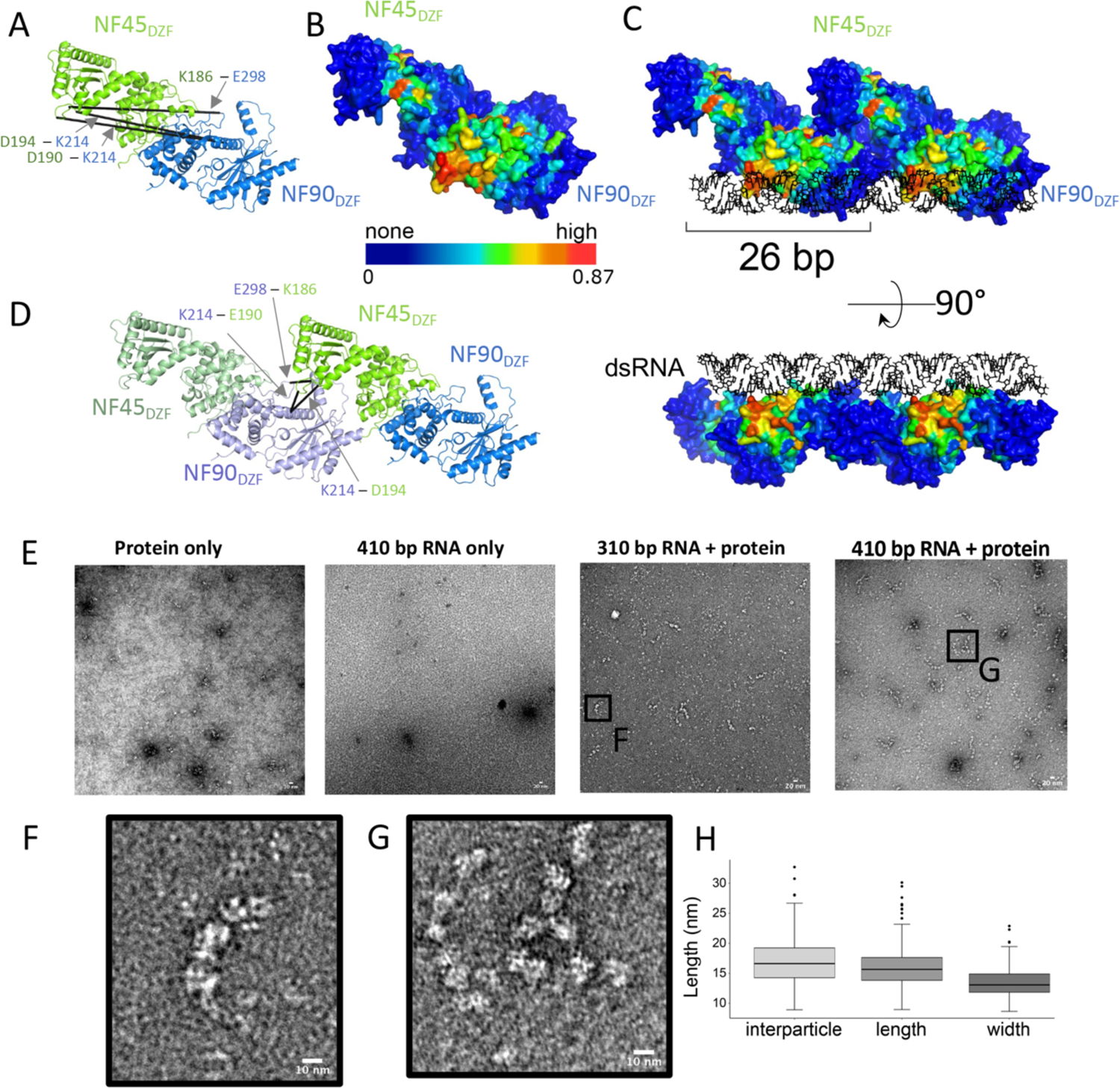
NF45-NF90_long_ complexes oligomerize on long stretches of dsRNA. (**A**) RNA-dependent cross-links between NF90_DZF_ and NF45_DZF_ displayed on a single heterodimer are longer than the distance constraint for EDC cross-linkers. (**B**) Surface of an NF45_DZF_-NF90_DZF_ heterodimer colored by likelihood of RNA binding, based on pyRBDome analyses (none-to-high, value range is given below gradient bar). (**C**) Combination of two NF45_DZF_-NF90_DZF_ heterodimers into a higher order oligomer model, showing a possible binding site for dsRNA. (**D**) Re-analysis of cross-links in (**A**) in the context of an open-ended oligomer model. (**E**) Negative stain electron micrograph of NF45-NF90_long_ protein alone; dsRNA with a maximum length of 410 bp; 10 mg/ml NF45-NF90_long_ mixed with dsRNA with a maximum length of 310 bp; 10 mg/ml NF45-NF90_long_ mixed with dsRNA with a maximum length of 410 bp. (**F**) Zoomed-in image of box in (**E**). (**G**) Zoomed-in images of box indicated in (**E**). (**H**) Quantitation of interparticle, length and width measurements from negative stain micrographs.

Lateral interactions between NF45-NF90 complexes could potentially provide an extended surface for binding dsRNA. While previous proteomic approaches identified peptides derived from DZF domains that are likely to directly contact RNA (Castello *et al*, 2016), we currently have no direct data to show where RNA binds on the surface of the DZF heterodimerisation domain. However, this structure has large areas of conserved residues that correlate with regions of positive charge (Fig. S5A). To predict likely RNA binding surfaces, we analysed NF45_DZF_-NF90_DZF_ models with pyRBDome (Chu *et al*., 2024), which generates ensemble predictions of protein-RNA interactions based on machine learning approaches. Models of the DZF heterodimerisation domain showed substantial areas of the protein with a high likelihood of RNA binding (Fig. 5B, Fig. S5A). Surfaces predicted to bind RNA overlap with areas of high conservation on the DZF heterodimerisation domain (Fig. 5B, S5A). We modelled the potential RNA binding surface by placing two NF45_DZF_-NF90_DZF_ complexes laterally; this generated a continuous binding surface with high likelihood of RNA binding (Fig. 5C, Fig. S5C). When queried against the cross-linking data, the excessively long cross-links within a DZF domain heterodimer (Fig. 5A) become short cross-links between heterodimers (Fig. 5D) and are well within the distance cut-off of the cross-linker. This lateral binding model provides good agreement with all the cross-linking data of the RNA-bound state (Fig. S5D). While the orientation of the RNA in this model is one of several possible poses, the footprint of a single NF45_DZF_-NF90_DZF_ heterodimer on the RNA approximates 26 bp (Fig. 5C, S5E). This agrees well with SAXS analysis (Fig. 3). Moreover, the model provides sufficient space for a second NF45-NF90 heterodimers to bind the opposite surface of the dsRNA, consistent with a 2:1 protein:RNA binding preference.

Collectively, the data and modelling suggest that NF45-NF90 complexes could form oligomers on extended lengths of dsRNA. To generate longer stretches of dsRNA, we used plasmids with opposing T7 promoters and varied the length of the transcribed sequences. Linearised plasmids were transcribed *in vitro* to generate dsRNA. RNA was then mixed with NF45-NF90_long_ complexes and examined by negative stain electron microscopy (EM) (Fig. 5E). We observed “beads on a string” structures in micrographs where protein-RNA complexes were characterised (Fig. 5E-G) that were absent from micrographs where only protein or RNA was stained (Fig. 5E). Measurement of the lengths and widths of these “beads” gave dimensions of ∼160-170 Å for interparticle distances and a similar length dimension for each particle, which is of the same magnitude as measured for single NF45-NF90_long_ in solution (Fig. 2E). Notably, the width of the particles averages around 130 Å (Fig. 5F). Measuring the width of the NF45_DZF_-NF90_DZF_ crystal structure gives dimensions of 45-65 Å. This suggests that the “beads” are likely to encompass two NF45-NF90_long_ complexes with dsRNA (20 Å diameter) sandwiched between them (Fig. 5H). The negative stain EM confirms that NF45-NF90_long_ complexes can form oligomeric structures that coat extended stretches of dsRNA.

## Discussion

The NF45-NF90 complex has many described roles in post-transcriptional regulation in mammals, including regulation of RNA A-to-I editing and control of cassette exon and circular RNA splicing (Freund *et al*., 2020; Haque *et al*., 2023; Li *et al*., 2017; Quinones-Valdez *et al*., 2019; Surka *et al*, 2021). In humans, the majority of NF45-NF90 is associated with inverted Alu repeats (Li *et al*., 2017; Quinones-Valdez *et al*., 2019; Van Nostrand *et al*., 2020) that are both the targets of A-to-I editing by ADAR1 (Bazak *et al*, 2014) and are highly enriched in introns (Zhang *et al*., 2014b). However, we still lack a molecular understanding of how this protein complex interacts with dsRNA and how such recognition events change the fates of individual RNA species. Prior work showed that dsRNA binding activity is dependent on the dsRBDs of NF90, yet the presence of the heterodimerised NF45_DZF_-NF90_DZF_ domains also contributes to RNA binding. Here we explore the impact of this larger domain on the architecture of the complex in solution without and with dsRNA.

Deep phylogenetic and sequence variant analysis showed that the DZF domains contribute substantially to NF45 and NF90 function (Fig. 1). Moreover, this analysis highlights several natively unstructured regions of NF110/NF90 that are conserved, indicating a contribution to function. While these contributions are not explored here, these two RGG motifs and a nearby RGRGRGRG sequence may contribute to RNA binding. The latter sequence is a substrate for PRMT5 symmetric methylation (Cox *et al*, 2022). Downstream of the RG repeat motif is a region of conservation and variant depletion shared in NF110 and NF90 isoforms (residues 660-680), suggestive of a short linear motif that could mediate protein-protein interactions. NF110 has further sites of localised conservation with benign missense variant depletion, around phosphorylation sites on residues 810, 812 and 816 and a further conserved motif at 845-850. These differences in motif distribution could reflect previously observed differences in sub-nuclear location and functions of NF110 and NF90 (Damianov *et al*., 2016; Viranaicken *et al*., 2011; Wandrey *et al*., 2015).

Extensive characterisation of the solution properties of NF45-NF90_long_ in comparison with two previously characterised fragments suggests that the longer complex is relatively compact in solution (Fig. S1, Fig. 2). Quantitative CLMS data indicates that this compact structure undergoes a substantial domain reorganisation on binding dsRNA. New interactions are observed between dsRBDs, as well as between individual dsRBDs and the DZF domains (Fig. 4). Notably, the conserved linker sequence between NF90_DZF_ and NF90_dsRBD1_ has strongly reduced crosslinking when dsRNA binds. This sequence overlaps with the nuclear localisation signal (residues 371-389), which may account for its conservation, yet it may also be an important contributor to domain reorganisation.

The binding of pairs of NF45-NF90_long_ complexes on dsRNA also creates a steric footprint on the dsRNA, such that around 26 bp are excluded by the binding event (Fig. 3). Analysis of RNA-dependent cross-links indicated that the heterodimerisation domain likely engages in lateral contacts when bound on dsRNA. This suggested that NF45-NF90 complexes could oligomerise on dsRNA, a model further supported by a composite prediction of likely RNA binding sites that fall primarily on NF90_DZF_ (Fig. 5). When lateral interactions are modelled, the outcome is a continuous, flat RNA binding surface that covers 26 bp of RNA, in good agreement with SAXS measurements (Fig. 3). Lateral interactions promote open-ended complexes or oligomers. We observed that NF45-NF90_long_ complexes can form “beads-on-a-string” structures on long dsRNAs (Fig. 5E,F). The dimensions of the “beads” suggest that they are pairs of complexes bound to dsRNA i.e. that they can coat the surface of the RNA (Fig. 5G).

That NF45-NF90 complexes can coat dsRNA suggests many possibilities of how NF45-NF90 complexes could regulate RNA fate. For example, in the context of A-to-I editing, coating of inverted Alu repeats could limit access of ADAR1 to its nuclear substrates, thus maintaining the low levels of editing observed at individual sites (Bazak *et al*., 2014). This would explain why loss of NF90 leads to a substantial increase in editing sites in human cells (Freund *et al*., 2020; Quinones-Valdez *et al*., 2019). In the context of cassette splicing, coating of long dsRNA structures by NF45-NF90 in introns could stabilise specific pre-mRNA secondary and tertiary structures. These stabilised sites could regulate access of the splicing machinery to splicing donor/acceptor sites, analogous to splicing regulation by polypyrimidine tract binding protein (Amir-Ahmady *et al*, 2005; Auweter & Allain, 2008; Liu *et al*, 2002). This would explain why loss of NF90 causes both alterations of splicing outcomes and splicing fidelity (Haque *et al*., 2023) and how NF90 promotes formation of circular RNAs (Li *et al*., 2017).

Coating of dsRNA by NF45-NF90 is reminiscent of other splicing regulators, such as heterogeneous nuclear ribonucleoprotein A1 (hnRNP A1), which can spread along ssRNA sequences from one high affinity site (Okunola & Krainer, 2009). Spreading of hnRNP A1 requires the presence of a glycine- and tyrosine-rich, intrinsically disordered region (IDR); similar IDRs in other hnRNP proteins were found to promote co-assemblies of these proteins on pre-mRNA (Gueroussov *et al*, 2017). While NF90 and NF110 contain similar IDRs at their C termini, these were not present in the constructs used in our analysis. It is possible that they could further promote the formation NF45-NF90 complexes on dsRNA, perhaps leading to cooperative effects, and this merits future study.

The interaction of NF45-NF90 DZF domains with dsRNA is also reminiscent of the interactions of OAS and cGAS with double-stranded nucleic acids (Civril *et al*., 2013; Donovan *et al*., 2013; Donovan *et al*., 2015; Hornung *et al*, 2014; Lohofener *et al*., 2015). Evolutionarily, cGAS and OAS are older than NF90, with homologs found in *Drosophila* and early metazoans (Cai & Imler, 2021; Wiens *et al*, 1999). In vertebrates, these enzymes recognise and respond to nucleic acids associated with viral infection. NF90, in its complex with NF45, may have emerged in vertebrates in response to a different kind of assault on cells, namely the expansion of elements like 7SL, B and Alu. By virtue of their high abundance, Alu elements contribute to extensive regions of dsRNA in the human transcriptome. NF45-NF90 complexes may have emerged to sequester these potentially problematic RNA structures in the nucleus. However, it is also possible that NF45-NF90 proteins have subsequently co-evolved to use AluIRs to shape co- and post-transcriptional events.

## Acknowledgements

We thank Diamond Light Source B21 staff for assistance with SAXS and mass photometry data collection. SAXS data are deposited in the SASDB, accession codes are SASDUA5, SASDUB5, SASDUC5, SASDUD5, SASDUE5, SASDUF5, SASDUG5, SASDUH5, SASDUJ5, SASDUK5, SASDUL5. MS data are deposited in PRIDE with accession PXD053010. This work utilised the Edinburgh Protein Production Facility (EPPF), the Centre Optical Instrumentation Laboratory funded by Wellcome Core Grants 092076 and 203149 to the Centre for Cell Biology. The authors would like to acknowledge Stephen Mitchell and Martin Singleton at the School of Biological Sciences Electron Microscopy unit for assistance with EM. We acknowledge the support of the Wellcome Multi-User Equipment Grant (WT104915MA). AGC and UJ were supported by a Wellcome Senior Fellowship (200898), SG was supported by Medical Research Council Senior Fellowship and Programme grants (MR/R008205/1 and MR/Y013131/1). SW was supported by BBSRC EASTBio doctoral training programme.

## Author contributions

SW – sample preparation, data collection and analysis, figure preparation, co-wrote manuscript

UJ – sample preparation, data collection and analysis, co-wrote manuscript

JZ – MS data collection, analysis and quantitation, co-wrote manuscript

JR – conceptualisation of quantitative MS, supervision, funding

SG – data analysis

AGC – conceptualisation of study, data collection and analysis, co-wrote manuscript, supervision, funding

## List of supplementary files

1. Supplementary Table 1 – DNA/RNA oligos used in this study
2. Supplementary Table 2 – SAXS experimental details
3. Supplementary Table 3 – quantified cross-links
4. Supplementary figures 1-5

