## Supplementary material for "Integrative structural analysis of NF45-NF90 heterodimers reveals architectural rearrangements and oligomerisation on binding dsRNA": Suppl. Table 1

Table 3 – Oligonucleotides used in this study

| **Name** | **RNA/**  **DNA/** | **Sequence 5’->3’** |
| --- | --- | --- |
| 18 bp top strand for EDC | RNA | UCACUUUCAUAAUGCUGG (Uniform 2’ Fluoro) |
| 18 bp bottom strand for EDC | RNA | CCAGCAUUAUGAAAGUGA |
| 25 bp GC-rich top strand for CLMS | RNA | GCCGCGGAGGCCCCGCCGUGGGCCC |
| 25 bp GC-rich bottom strand for CLMS | RNA | GGGCCCACGGCGGGGCCUCCGCGGC |
| 25 bp for SAXS top strand | RNA | UCACUUUCAUAAUGCUGGUCACUUU |
| 25 bp for SAXS bottom strand | RNA | AUGCUGGUCACUUUCAUAAUGCUGG |
| 36 bp for SAXS top strand | RNA | UCACUUUCAUAAUGCUGGUCACUUUCAUAAUGCUGG |
| 36 bp for SAXS bottom strand | RNA | CCAGCAUUAUGAAAGUGACCAGCAUUAUGAAAGUGA |
| 54 bp for SAXS top strand | RNA | UCACUUUCAUAAUGCUGGUCACUUUCAUAAUGCUGGUCACUUUCAUAAUGCUGG |
| 54 bp for SAXS bottom strand | RNA | CCAGCAUUAUGAAAGUGACCAGCAUUAUGAAAGUGACCAGCAUUAUGAAAGUGA |
| **PCR primers** |  |  |
| 310bp_Cyp1A1_F | DNA | 5’ P aatgccgttttattccgatttc |
| 310bp_Cyp1A1_R | DNA | 5’ P gttattgaagttcccggacac |
