## Supplementary material for "Integrative structural analysis of NF45-NF90 heterodimers reveals architectural rearrangements and oligomerisation on binding dsRNA": Suppl. Table 2

Table 2. SAXS experimental details and data parameters for NF45-NF90_long_ with dsRNA

| (a) Sample details | | | | | | | | | | | | |
| --- | --- | --- | --- | --- | --- | --- | --- | --- | --- | --- | --- | --- |
|  | Organism | | Source | | | UniProt ID (residues in construct) + uncleaved tag | | | | | Molecular mass (Da) | |
| NF45 | *Homo sapiens* | | *E.coli* expressed | | | Q12905 (1-390) + GST-tag (GPLGSPEF) at N-terminus | | | | | 43846 | |
| NF90 | *Mus musculus* | | *E. coli* expressed | | | Q9Z1X4 (1-591) + Additional G + LE scar site with HHHHHH-tag at C-terminus | | | | | 65897 | |
| 25mer | - | | Biomers RNA oligos | | | 5’ UCA CUU UCA UAA UGC UGG UCA CUU U 3’ 3’ CCA GCA UUA UGA AAG UGA CCA GCA U 5’ | | | | | 15949 | |
| 36mer | - | | Biomers RNA oligos | | | 5’ UCA CUU UCA UAA UGC UGG UCA CUU UCA UAA UGC UGG 3’  3’ CCA GCA UUA UGA AAG UGA CCA GCA UUA UGA AAG UGA 5’ | | | | | 23102 | |
| 54mer | - | | Biomers RNA oligos | | | 5’ UCA CUU UCA UAA UGC UGG UCA CUU UCA UAA UGC UGG UCA CUU UCA UAA UGC UGG 3’ 3’ CCA GCA UUA UGA AAG UGA CCA GCA UUA UGA AAG UGA CCA GCA UUA UGA AAG UGA 5’ | | | | | 65897 | |
| Samples | | NF90-NF45 | NF90-NF45 25mer 2:1 | | NF90-NF45 25mer 4:1 | | NF90-NF45 36mer 2:1 | NF90-NF45 36mer 4:1 | NF90-NF45 54mer 2:1 | NF90-NF45 54mer 4:1 | | NF90-NF45 54mer 6:1 |
| Column | | | | SEC-SAXS column, s200 increase 3.2/200 | | | | | | | | |
| Loading concentration (mg/ml) | | | | 5.5 | | | | | | | | |
| Injection volume (µl) | | | | 60 | | | | | | | | |
| Flow rate (ml/min) | | | | 0.075 | | | | | | | | |
| Solvent composition | | | | 20 mM HEPES pH 7.5 150 mM NaCl 1 mM DTT | | | | | | | | |

| (b) SAC-SAXS data collection | |
| --- | --- |
| Instrument | Diamond Light Source Ltd Synchrotron, BL21 beamline, EigerX 4M detector (Dectris) |
| Source | Bending magnet |
| Wavelength (Å) | 0.9464 |
| Beam size at focal point (µm) | 50×50 |
| Sample-to-detector distance (m) | 3.7193 |
| q-measurement range (Å^-1^) | 0.0045-0.34 |
| Exposure time | Continuous 0.005 s data-frame measurements of SEC elution. |
| Frames | 600 |
| Sample temperature (°C) | 15 |

| (c) Software employed for SAS data reduction, analysis and interpretation | |
| --- | --- |
| Sample – Solvent subtraction | Chromixs (Panjkovich and Svergun, 2018) from ATSAS 3.2.1 (Manalastas-Cantos et al., 2021) |
| Basic analyses: Guinier, P(r), V_p_ | ScÅtter IV (https://bl1231.als.lbl.gov/scatter/) |
| Shape/bead modelling | DAMMIF (Franke and Svergun, 2009) and DAMMIN (Svergun, 1999) via ATSAS 3.2.1 (Manalastas-Cantos et al., 2021) |
| Atomic structure modelling | MultiFoXS (Schneidman-Duhovny et al., 2016) via web server (https://modbase.compbio.ucsf.edu/multifoxs/) |
| Molecular graphics | PyMOL 2.5.4 |

| (d) Structural parameters | | | | | | | | |
| --- | --- | --- | --- | --- | --- | --- | --- | --- |
|  | NF90-NF45 | NF90-NF45 25mer 2:1 | NF90-NF45 25mer 4:1 | NF90-NF45 36mer 2:1 | NF90-NF45 36mer 4:1 | NF90-NF45 54mer 2:1 | NF90-NF45 54mer 4:1 | NF90-NF45 54mer 6:1 |
| I(0) (cm^-1^) [from Guinier] | 0.089 | 0.1905 | 0.08928 | 0.2563 | 0.2354 | 0.3006 | 0.4032 | 0.229 |
| R_g_ (Å) [from Guinier] | 48.03 ± 0.08089 | 58.95 ± 0.06254 | 58.12 ± 0.08977 | 63.90 ± 0.09438 | 63.62 ± 0.09487 | 73.86 ± 0.1347 | 79.29 ± 0.08955 | 79.47 ± 0.1291 |
| q_min_R_g_ - q_max_R_g_ used for Guinier | 0.6519 - 1.2150 | 0.8073 - 1.2314 | 0.6406 - 1.1960 | 0.8198 - 1.2037 | 0.7086 - 1.1893 | 1.0175 - 1.2406 | 0.7101 - 1.2339 | 0.7132 - 1.1685 |
| Score | 1.023 | 1.087 | 1.03 | 1.075 | 1.076 | 1.073 | 1.069 | 1.079 |
| I(0) (cm^-1^) [from p(r)] | 0.0801 | 0.1714 | 0.08193 | 0.2324 | 0.2157 | 0.2676 | 0.3579 | 0.2069 |
| R_g_ (Å) [from p(r)] | 46.77 | 57.05 | 56.9 | 61.57 | 63.73 | 71.63 | 77.66 | 77.54 |
| D_max_ (Å) [from p(r)] | 171 | 198.5 | 204.5 | 212 | 220.5 | 250 | 264.5 | 260.5 |
| Porod volume, V_p_ (Å^-3^) | 234456 | 398098 | 399715 | 587120 | 595567 | 613121 | 788402 | 791908 |
| Volume-of-correlation, V_c_ | 813.3 | 1278.9 | 1239.1 | 1461.2 | 1532.9 | 1759.4 | 2108.4 | 2009 |
| Molecular mass (kDa) [from V_p_] | 141.2 | 239.8 | 240.8 | 353.7 | 358.8 | 369.4 | 474.9 | 477.1 |

| (e) Shape modelling results | | | | | | | | |
| --- | --- | --- | --- | --- | --- | --- | --- | --- |
| DAMMIF (default parameters, 10 repetitions), averaged with DAMAVER and refined with DAMMIN | | | | | | | | |
|  | NF90-NF45 | NF90-NF45 25mer 2:1 | NF90-NF45 25mer 4:1 | NF90-NF45 36mer 2:1 | NF90-NF45 36mer 4:1 | NF90-NF45 54mer 2:1 | NF90-NF45 54mer 4:1 | NF90-NF45 54mer 6:1 |
| Symmetry/ anisotropy assumption | P1, none | P1, none | P1, none | P1, none | P1, none | P1, none | P1, none | P1, none |
| χ^2^ value | 1.274 | 1.4 | 1.067 | 1.309 | 1.354 | 1.288 | 1.201 | 1.491 |
| Constant subtraction procedure | 3.37×10^-4^ | 6.60×10^-4^ | 2.93×10^-4^ | 1.09×10^-3^ | 1.01×10^-3^ | 1.96×10^-3^ | 2.72×10^-3^ | 1.50×10^-3^ |
| Model resolution (SASRES) (Å) | 44.4151 | 56.0733 | 45.6861 | 61.2184 | 60.5901 | 69.8233 | 76.7173 | 68.5343 |

| (f) Atomistic modelling | NF90-NF45 |
| --- | --- |
| Multistate/ensemble models | MultiFoXS (10 000 models in starting set) |
| Starting crystal structures | 4AT7, 5DV7, AF-Q9Z1X4, AF-Q9CXY6 |
| Flexible residues | 1-28A, 362-390A, 1-5B, 55-86B, 341-353B, 376-381B, 468-518B, 591B |
| No. of states | 5 |
| χ^2^ CORMAP p-values | 1.23 |
| c1, c2 | 1.02, 0.31 |
| Rg values of each state (Å) | 1) 50.76, 2) 43.81, 3) 58.09, 4) 52.30, 5) 39.54 |
| Weights wn | w1: 0.224, w2: 0.304, w3: 0.172, w4: 0.15, w5: 0.15 |

| (g) Data and model deposition IDs | | | | | | | | |
| --- | --- | --- | --- | --- | --- | --- | --- | --- |
|  | NF90-NF45 | NF90-NF45 25mer 2:1 | NF90-NF45 25mer 4:1 | NF90-NF45 36mer 2:1 | NF90-NF45 36mer 4:1 | NF90-NF45 54mer 2:1 | NF90-NF45 54mer 4:1 | NF90-NF45 54mer 6:1 |
| SASBDB | SASDUC5 | SASDUE5 | SASDUF5 | SASDUG5 | SASDUH5 | SASDUJ5 | SASDUK5 | SASDUL5 |

Table 2. SAXS experimental details and data parameters for NF90-NF45 constructs

| (a) Sample details | | | | | |
| --- | --- | --- | --- | --- | --- |
| Sample | NF90_long_-NF45 | | NF90_DZF_-NF45_DZF_ | | NF90_dsRBDs_ |
|  | NF90 | NF45 | NF90 | NF45 | NF90 |
| Organism | *Mus musculus* | *Homo sapiens* | *Mus musculus* | *Mus musculus* | *Mus musculus* |
| Source (Catalogue No. or reference) | *E. coli* expressed | *E. coli* expressed | *E. coli* expressed | *E. coli* expressed | *E. coli* expressed |
| Uniprot ID (residues in construct) + uncleaved tag | Q9Z1X4 (1-591) + Additional G + LE scar site + HHHHHH-tag at C-terminus | Q12905 (1-390) + GST-tag (GPLGSPEF) at N-terminus | Q9Z1X4 1-381 + GH TEV cleavage scar at N-terminus | Q9CXY6 29-390 + GS at N-terminus from precision site | Q9Z1X4 395-592 + Additional G at N-terminus from GST-tag |
| Molecular mass M from chemical composition (Da) | 65897 | 43846 | 42468 | 40320 | 21138 |
| Column | SEC-SAXS column, s200 increase 3.2/200 | | | | |
| Loading concentration (mg/ml) | 7.75 | | 10 | | 10 |
| Injection volume (µl) | 60 | | 60 | | 60 |
| Flow rate (ml/min) | 0.1 | | 0.1 | | 0.1 |
| Solvent composition | 20 mM HEPES pH 7.5 150 mM NaCl 1 mM DTT | | | | |

| (b) SAS data collect parameters | |
| --- | --- |
| Instrument | Diamond Light Source Ltd Synchrotron, BL21 beamline, EigerX 4M detector (Dectris) |
| Source | Bending magnet |
| Wavelength (Å) | 0.954 |
| Beam size at focal point (µm) |  |
| Sample-to-detector distance (m) | 3.7 |
| q-measurement range (Å^-1^) |  |
| Exposure time | Continuous 0.005 s data-frame measurements of SEC elution (915 frames) |
| Sample temperature (°C) | 15 |

| (c) Software employed for SAS data reduction, analysis and interpretation | |
| --- | --- |
| Sample – Solvent subtraction | ScÅtter IV (https://bl1231.als.lbl.gov/scatter/) |
| Basic analyses: Guinier, P(r), V_p_ | ScÅtter IV (https://bl1231.als.lbl.gov/scatter/) |
| Shape/bead modelling | GASBOR 2.3i (Svergun et al., 2001) |
| Atomic structure modelling | MultiFoXS (Schneidman-Duhovny et al., 2016) via web server (https://modbase.compbio.ucsf.edu/multifoxs/) |
| Molecular graphics | PyMOL 2.5.4 |

| (d) Structural parameters | | | |
| --- | --- | --- | --- |
|  | NF90_long_-NF45 | NF90_DZF_-NF45_DZF_ | NF90_dsRBDs_ |
| I(0) (cm^-1^) [from Guinier] | 0.0894 | 0.119 | 0.0346 |
| R_g_ (Å) [from Guinier] | 42.53 ± 0.177 | 36.08 ± 0.6563 | 35.82 ± 0.8592 |
| qminRg - qmaxRg used for Guinier | 0.6692 - 1.1881 | 0.7444 - 1.2842 | 0.7024 - 1.0828 |
| Score | 1.183 | 1.076 | 1.043 |
| I(0) (cm^-1^) [from p(r)] | 0.101 | 0.119 | 0.0309 |
| R_g_ (Å) [from p(r)] | 46.93 | 35.93 | 34.68 |
| D_max_ (Å) [from p(r)] | 171 | 125.5 | 126 |
| Porod volume, Vp (Å^-3^) | 270388 | 167777 | 48622 |
| Volume-of-correlation, Vc | 821.9 | 590.5 | 314.1 |
| Molecular mass (kDa) [from Vp] | 162.9 | 101.1 | 29.3 |
| Molecular mass (kDa) [from Vc] | 116.8 | 80.2 | 22.8 |

| (e) Shape modelling results | | | |
| --- | --- | --- | --- |
|  | NF90_long_-NF45 | NF90_DZF_-NF45_DZF_ | NF90_dsRBDs_ |
| GASBOR (default parameters) |  |  |  |
| Number of dummy residues | 998 | 747 | 199 |
| Symmetry/anisotropy assumptions | P1, none | P1, none | P1, none |
| χ^2^ value | 1.413 | 1.079 | 1.301 |

| (f) Atomistic modelling | | | |
| --- | --- | --- | --- |
| Multistate/ensemble models | | | |
| MultiFoXS (10 000 models in starting set) | | | |
|  | NF90_long_-NF45 | NF90_DZF_-NF45_DZF_ | NF90_dsRBDs_ |
| Starting crystal structures | 4AT7, 5DV7, AF-Q9Z1X4, AF-Q9CXY6 | 4AT7, 5DV7, AF-Q9Z1X4, AF-Q9CXY6 | 4AT7, 5DV7, AF-Q9Z1X4, AF-Q9CXY6 |
| Flexible residues | 1-28A, 362-390A, 1-5B, 55-86B, 341-353B, 376-381B, 468-518B, 591B | 362-390A, 1-5B, 55-86B, 341-353B, 376-381B | 394-402A, 468-518A |
| No. of states | 4 | 1 | 4 |
| χ^2^ CORMAP p-values | 1.18 | 1.23 | 1.04 |
| c1, c2 | 1.00, 1.03 | 1.02, 0.35 | 1.03, 1.80 |
| Rg values of each state (Å) | 1) 46.74 | 1) 35.47 | 1) 31.27 |
|  | 2) 47.97 |  | 2) 42.23 |
|  | 3) 54.78 |  | 3) 56.66 |
|  | 4) 39.51 |  | 4) 28.59 |
| Weights wn | w1: 0.095 | w1: 1.0 | w1: 0.553 |
|  | w2: 0.587 |  | w2: 0.314 |
|  | w3: 0.246 |  | w3: 0.08 |
|  | w4: 0.072 |  | w4: 0.052 |

| (g) Data and model deposition IDs | | | |
| --- | --- | --- | --- |
|  | NF90_long_-NF45 | NF90_DZF_-NF45_DZF_ | NF90_dsRBDs_ |
| SASBDB | SASDUD5 | SASDUB5 | SASDUA5 |

FRANKE, D. & SVERGUN, D. I. 2009. DAMMIF, a program for rapid ab-initio shape determination in small-angle scattering. *J Appl Crystallogr,* 42**,** 342-346.

MANALASTAS-CANTOS, K., KONAREV, P. V., HAJIZADEH, N. R., KIKHNEY, A. G., PETOUKHOV, M. V., MOLODENSKIY, D. S., PANJKOVICH, A., MERTENS, H. D. T., GRUZINOV, A., BORGES, C., JEFFRIES, C. M., SVERGUN, D. I. & FRANKE, D. 2021. ATSAS 3.0: expanded functionality and new tools for small-angle scattering data analysis. *J Appl Crystallogr,* 54**,** 343-355.

PANJKOVICH, A. & SVERGUN, D. I. 2018. CHROMIXS: automatic and interactive analysis of chromatography-coupled small-angle X-ray scattering data. *Bioinformatics,* 34**,** 1944-1946.

SCHNEIDMAN-DUHOVNY, D., HAMMEL, M., TAINER, J. A. & SALI, A. 2016. FoXS, FoXSDock and MultiFoXS: Single-state and multi-state structural modeling of proteins and their complexes based on SAXS profiles. *Nucleic Acids Research,* 44**,** W424-W429.

SVERGUN, D. I. 1999. Restoring low resolution structure of biological macromolecules from solution scattering using simulated annealing. *Biophys J,* 76**,** 2879-86.

SVERGUN, D. I., PETOUKHOV, M. V. & KOCH, M. H. 2001. Determination of domain structure of proteins from X-ray solution scattering. *Biophys J,* 80**,** 2946-53.
