## Supplementary Figures for "Integrative structural analysis of NF45-NF90 heterodimers reveals architectural rearrangements and oligomerisation on binding dsRNA"

### Figure S1

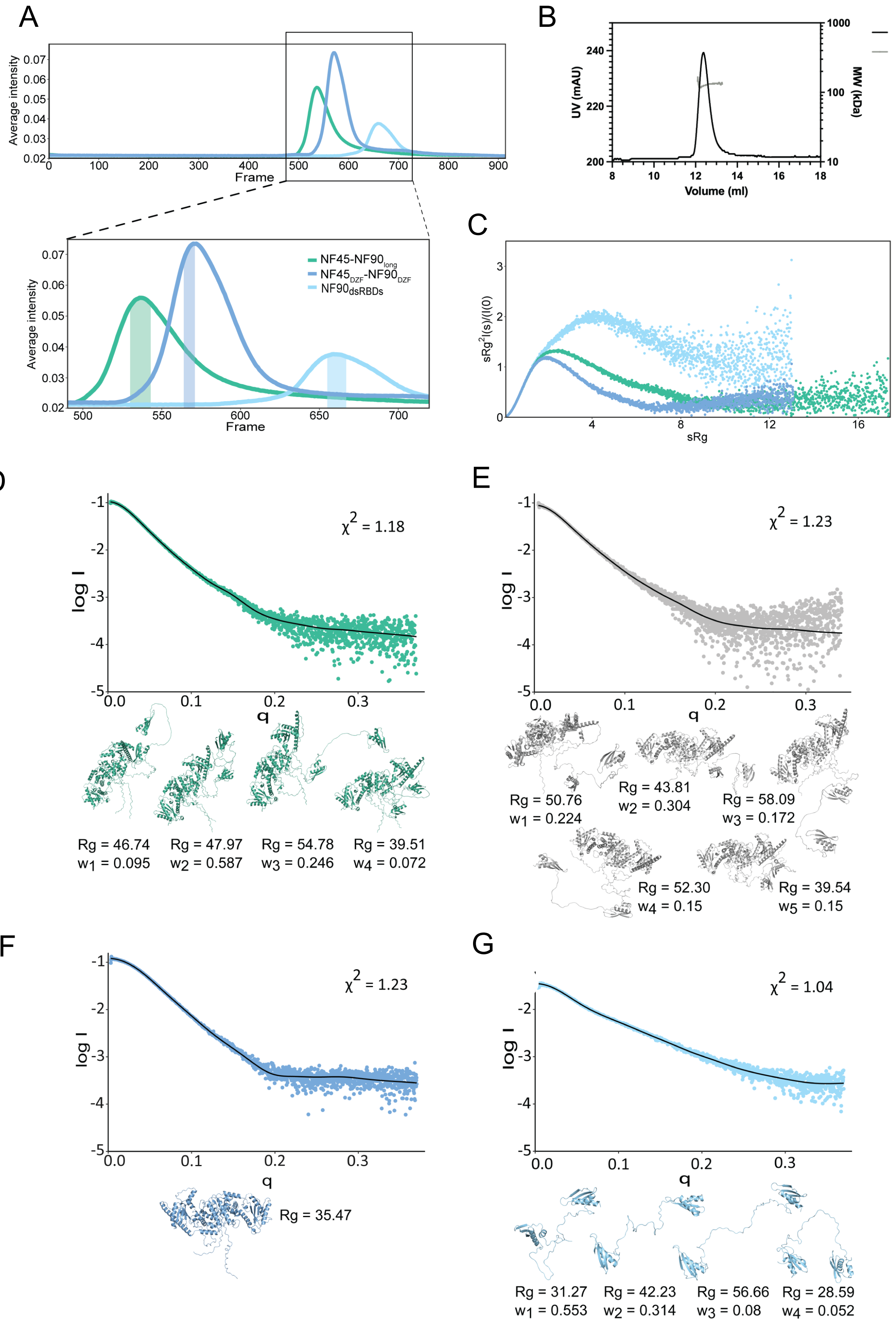


**Figure S1**. NF45-NF90 complexes in solution show compaction of domains. (**A**) SEC-SAXS profiles of three constructs with inset showing the data used for subsequent analysis. (**B**) SEC-MALS analysis of NF45-NF90$_{long}$. (**C**) Normalised Kratky plots for the three constructs analysed in (**A**). (**D-G**) Conformational diversity of NF45-NF90 is consistent with SAXS data. The fitting to SAXS data is given by the black line in each case. MultiFoXS analysis with (**D**) and (**E**) NF45-NF90$_{long}$ from two independent datasets, (**F**) NF45$_{DZF}$-NF90$_{DZF}$ with a single model, (**G**) NF90$_{dsRBDs}$ with four models.

### Figure S2

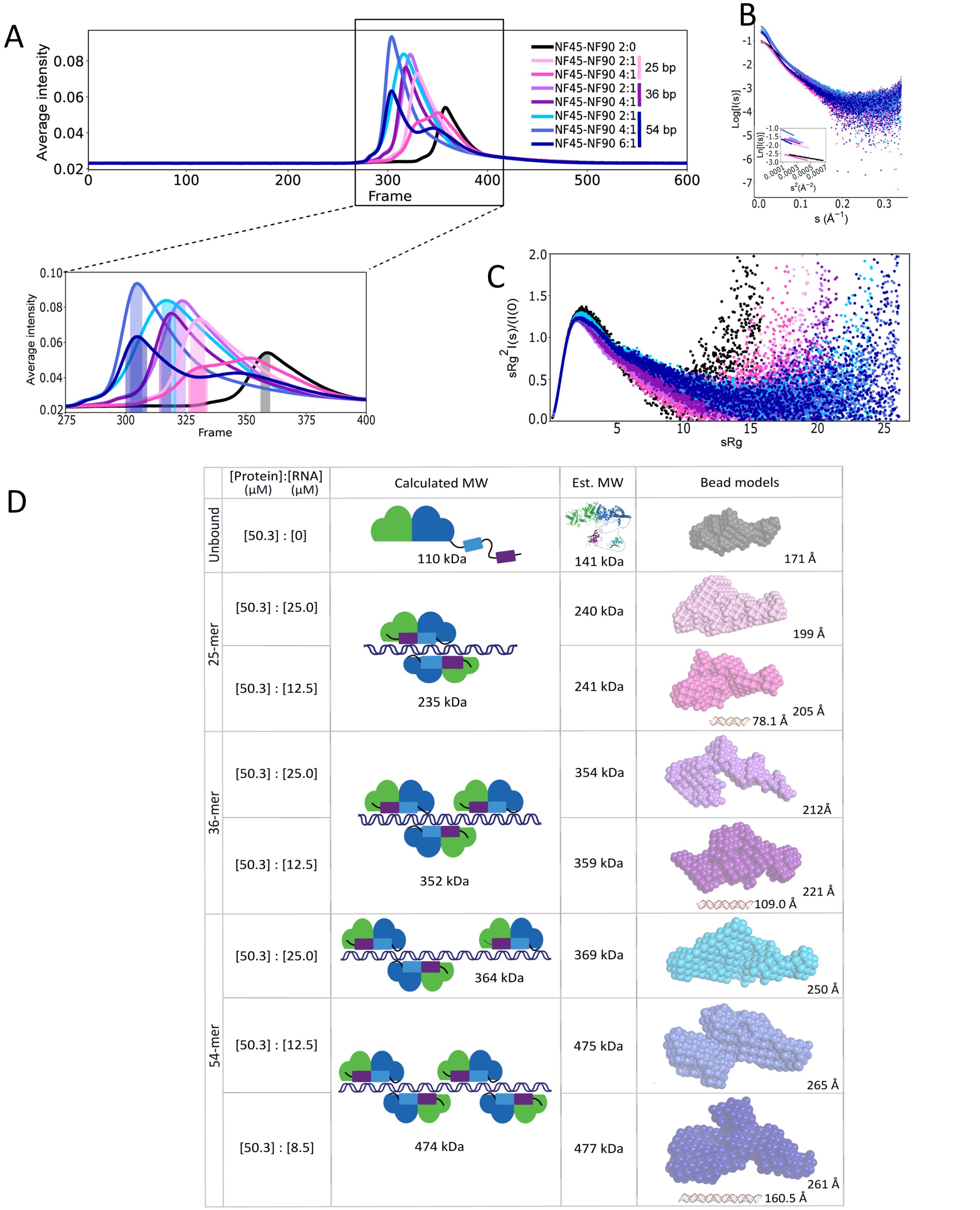


**Figure S2**. Solution analysis of NF45-NF90 binding to dsRNA of increasing lengths. Eight SAXS measurements were carried out with no dsRNA or with 25 bp, 36 bp and 54 bp RNA at increasing molar ratios of protein:dsRNA. (**A**) Intensity profiles of samples as eluted from size exclusion chromatography (Fig. 4A) with zoomed inset figures showing frames selected for real space analysis. (**B**) Extracted scattering curves for all samples with inset of Guinier analysis. (**C**) Normalised Kratky plots for all SAXS samples. (**D**) A summary of SAXS analysis. Molar ratios of complexes are shown in the first column. Likely molecular compositions with calculated molecular masses are shows as cartoon models in the centre, with SAXS-derived masses and bead models shown on the right as in Fig. 3B. $D_{max}$ values are indicated under each model with an RNA model to scale.

### Figure S3

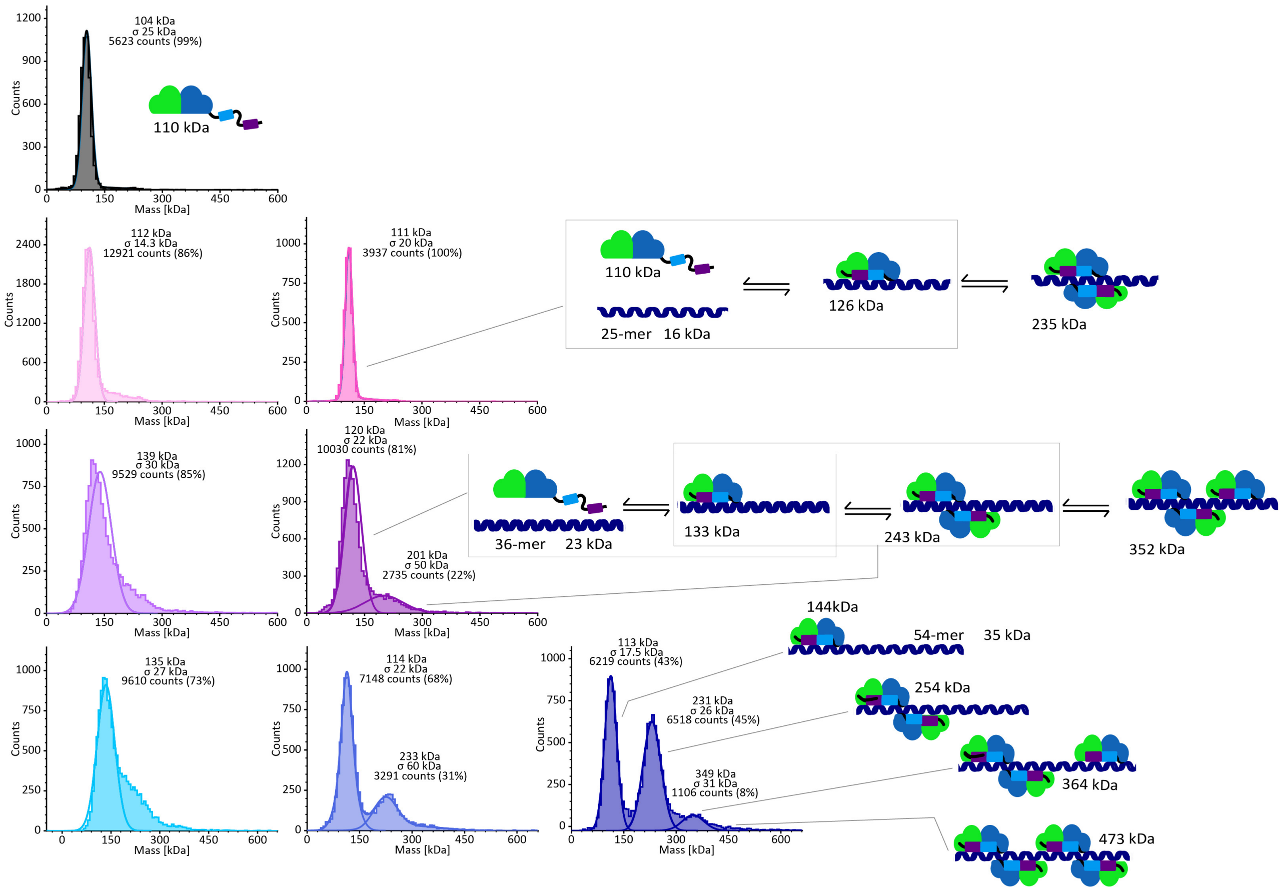


**Figure S3:** Population mass distributions of NF45-NF90$_{long}$ with increasing lengths of dsRNA by mass photometry. Masses given above peaks are median values, with associated standard deviations (σ), providing estimates of the molecular weight. Grey: NF45-NF90 complexes alone (109 kDa). Pink: NF45-NF90 in a 2:1 ratio and 4:1 ratio with 25 bp dsRNA; Purple: NF45-NF90 in a 2:1 and 4:1 ratio with 36 bp dsRNA; Blue: NF45-NF90 in a 2:1, 4:1 or 6:1 ratio with 54 bp dsRNA. Cartoons depict the likely equilibria and species present in each sample, with calculated molecular weights for those species.

### Figure S4

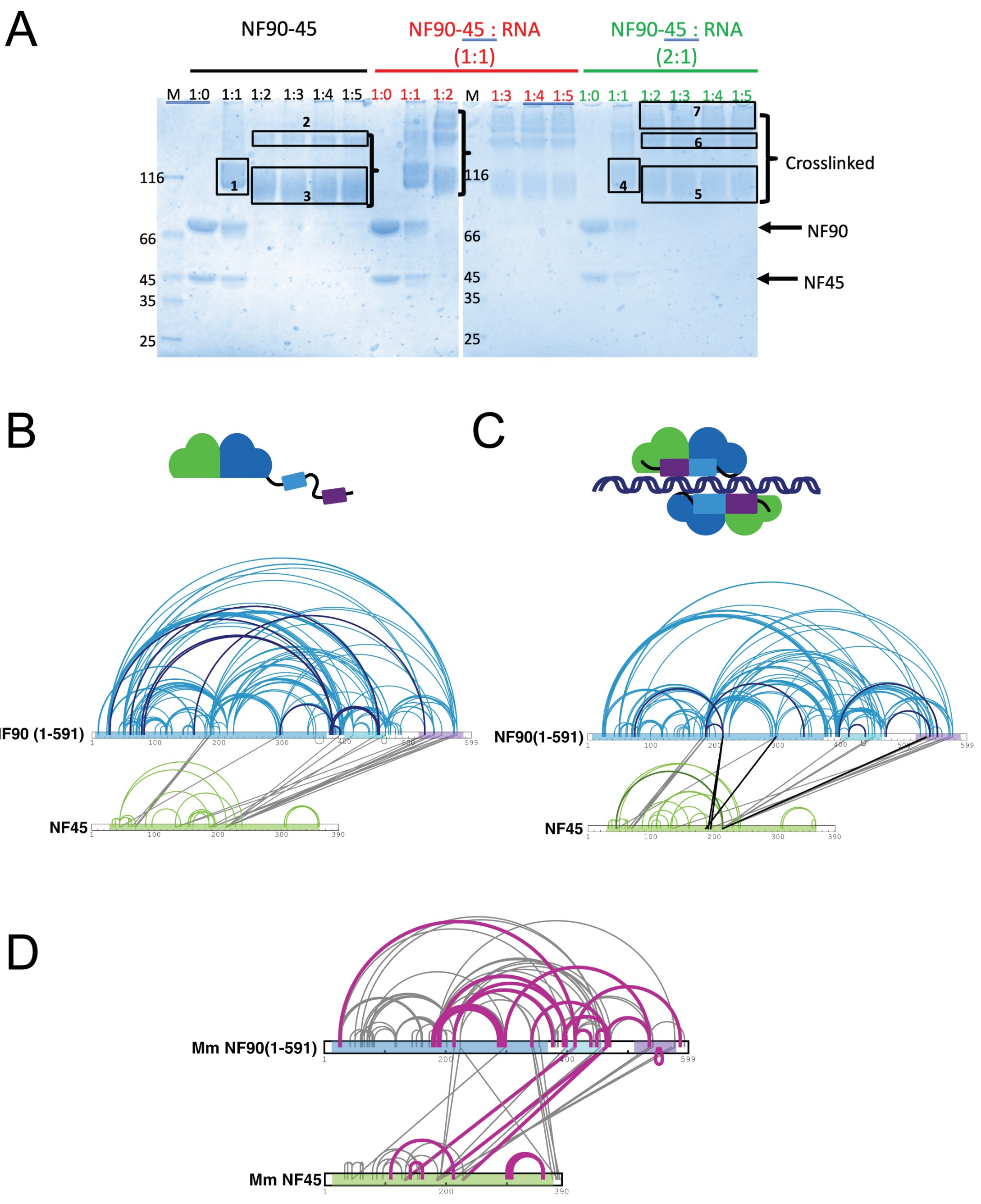


**Figure S4**. EDC crosslinking of NF45-NF90$_{long}$ complexes. (**A**) SDS-PAGE analysis of cross-linked samples, showing titration of EDC crosslinker with different samples of NF45-NF90$_{long}$. Boxes indicate the samples that were included in analysis. Samples 1-3 were used for data presented in Fig. 4A; Samples 4-6 were used for data present in Fig. 4B. (**B**) Data corresponding to Fig. 4C showing both quantified (dark) and non-quantified cross-links from this dataset. (**C**) Data corresponding to Fig. 4D showing both quantified (dark) and non-quantified cross-links from this dataset. (**D**) Data from all RNA bound samples including sample 7, showing cross-links specific to sample 7 in purple.

**Figure S5**

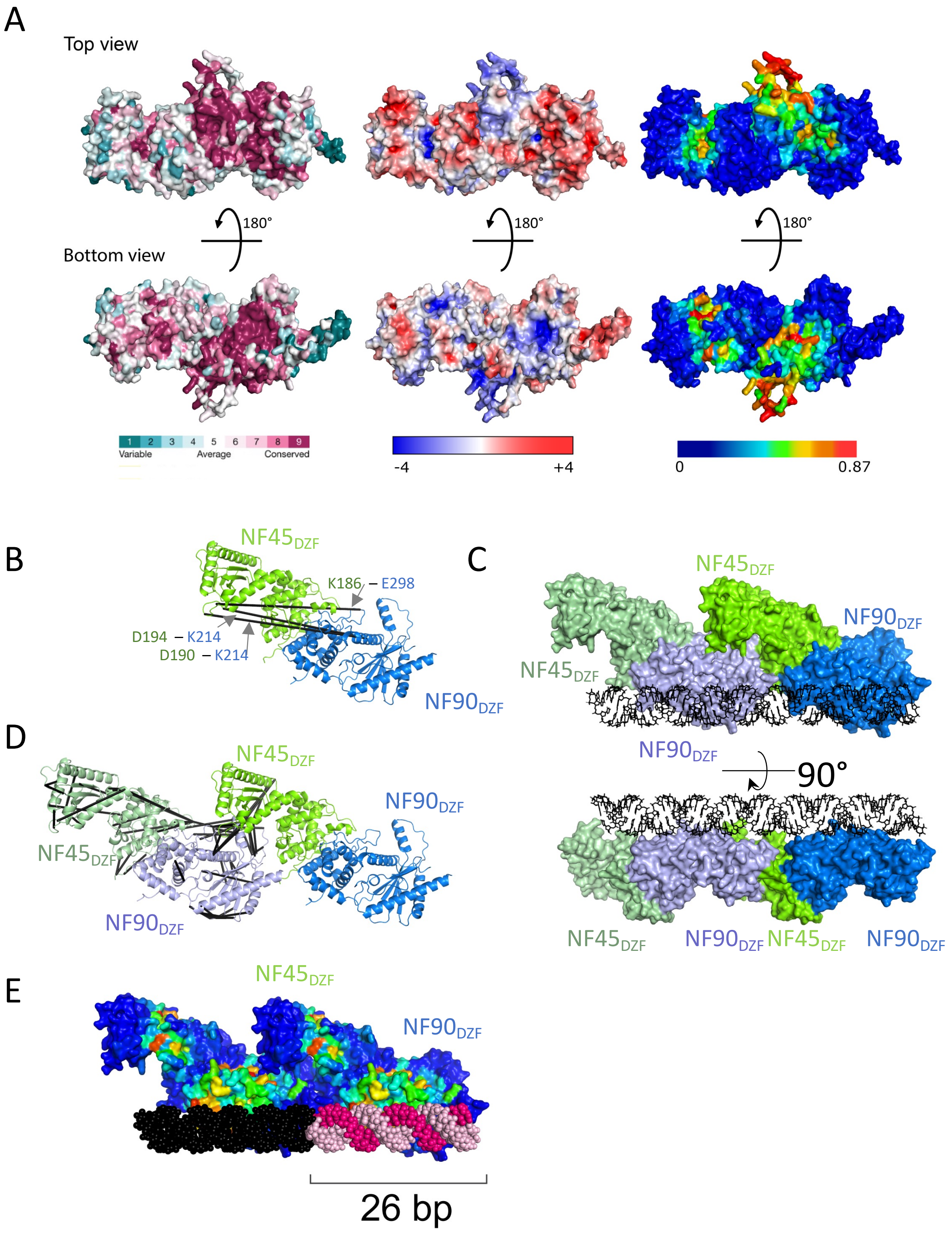


**Figure S5**. NF45-NF90$_{long}$ complexes oligomerize on long stretches of dsRNA. (**A**) Comparison of surface characteristics of NF45$_{DZF}$-NF90$_{DZF}$ showing conservation, electrostatics (-4 to +4 kTe) and RNA binding propensity (none to high). (**B**) NF45$_{DZF}$-NF90$_{DZF}$ heterodimer as shown in Fig. 5A, reproduced here for comparison with (**D**). Crosslinks are shown as black lines. (**C**) Similar model to Fig. 5C but with surfaces colored by molecule. (**D**) Re-analysis of cross-links in (**B**) in the context of an open-ended oligomer model and displaying all measured cross-links. (**E**) Model as in (**C**) but highlighting RNA binding propensity and showing the dsRNA model as black or pink to indicate the length of 26 bp.
